# Contour junctions underlie neural representations of scene categories in high-level human visual cortex: Contour junctions underlie neural code of scenes

**DOI:** 10.1101/044156

**Authors:** Heeyoung Choo, Dirk. B. Walther

**Affiliations:** Department of Psychology, University of Toronto 100 St. George Street, Toronto, ON, M5S 3G3 Canada

**Keywords:** Neural representations of scenes, encoding of scene structure, contour junctions, scene categorization, parahippocampal place area, occipital place area, visual cortex, fMRI, multi-voxel pattern analysis

## Abstract

Humans efficiently grasp complex visual environments, making highly consistent judgments of entry-level category despite their high variability in visual appearance. How does the human brain arrive at the invariant neural representations underlying categorization of real-world environments? We here show that the neural representation of visual environments in scenes-selective human visual cortex relies on statistics of contour junctions, which provide cues for the three-dimensional arrangement of surfaces in a scene. We manipulated line drawings of real-world environments such that statistics of contour orientations or junctions were disrupted. Manipulated and intact line drawings were presented to participants in an fMRI experiment. Scene categories were decoded from neural activity patterns in the parahippocampal place area (PPA), the occipital place area (OPA) and other visual brain regions. Disruption of junctions but not orientations led to a drastic decrease in decoding accuracy in the PPA and OPA, indicating the reliance of these areas on intact junction statistics. Accuracy of decoding from early visual cortex, on the other hand, was unaffected by either image manipulation. We further show that the correlation of error patterns between decoding from the scene-selective brain areas and behavioral experiments is contingent on intact contour junctions. Finally, a searchlight analysis exposes the reliance of visually active brain regions on different sets of contour properties. Statistics of contour length and curvature dominate neural representations of scene categories in early visual areas and contour junctions in high-level scene-selective brain regions.

## 1. Introduction

When humans view their complex natural environment, they have rapid access to several aspects of its content, such as identity and category of the scene, presence of particular object categories, or global layout (Potter & Levy, 1969; Thorpe, Fize, & Marlot, 1996; Fei-Fei, Iyer, Koch, & Perona, 2007; Greene & Oliva, 2009a, 2009b, 2010). We here investigate the neural mechanisms for the detection of cues to the three-dimensional structure of complex real-world scenes in human visual cortex: We show that the neural representation of scene categories in several high-level visual brain regions, but not in early visual cortex, critically depends on contour junctions.

Entry-level category of a scene is a central aspect of the human visual perception of real-world environment (Tversky & Hemenway 1983). Recent evidence suggests that humans compulsively categorize scenes even if it is detrimental to their task (Greene & Fei-Fei, 2014). Scene categories can be decoded from the parahippocampal place area (PPA) of humans passively viewing scene images (Walther, Caddigan, Fei-Fei, & Beck, 2009; Park, Bradly, Greene, & Oliva, 2011; Walther, Chai, Caddigan, Fei-Fei, & Beck, 2011). Moreover, error patterns for category decoding from the PPA match the pattern of human errors during a rapid scene categorization task (Walther et al., 2009).

There has been considerable debate over the visual properties that underlie the neural representation of scene categories. According to one popular hypothesis, statistics of orientations at different scales as captured by the Fourier amplitude spectrum make accurate computational predictions about entry-level categories of real-world scene images. Subsequent principal component analysis revealed that several diagnostic structures in the Fourier amplitude spectrum are directly related to global scene properties, such as openness, naturalness, or mean distance (Oliva & Torralba, 2001; Torralba & Oliva, 2003). These global properties, in turn, are thought to give rise to a representation of scene categories (Greene & Oliva, 2009a, 2009b, 2010). For instance, the “beach” category is represented as an open natural environment whereas the highway category is represented as an open and man-made environment. In support of this hypothesis, several global scene properties have been found to be represented in activity patterns in the PPA (Harel, Kravitz, & Baker, 2013; Kravitz, Peng, & Baker, 2011; Park et al., 2011, Park, Konkle, & Oliva, 2015).

Here, we posit that real-world scene categories are built by recovering the three-dimensional shape of the visual world rather than relying on orientation statistics. The visual world can be described by the relations of surfaces and shapes in space (e.g., by the 2½-dimensional sketch of a scene; Marr, 1982). Contour junctions available in the two-dimensional scene images diagnostically describe these spatial relations. For instance, L-junctions indicates points of termination of surfaces, T-junctions signify occlusion in depth, and Y- and arrow-junctions indicate corners facing toward or away from the viewer. Angles of contour junctions indicate the extent to which depth changes over surfaces (Biederman, 1987). The diagnostic value of contour junction properties holds for simple artificial scenes consisting of geometric objects (Guzman, 1968) as well as for real-world object recognition (Biederman, 1987). Furthermore, a computational model based on category-specific statistics of contour junction properties explained human errors in rapid categorization of real-world scene images (Walther & Shen, 2014). According to this structural representation hypothesis, contour junctions should be tied to the neural representation of complex real-world scenes.

Line drawings are a powerful tool to investigate scene recognition, even though they are impoverished depictions of scenes, compared to full-textured color photographs. In fact, line drawings can be categorized or recognized as quickly and accurately as full-textured color photographs (Biederman & Ju, 1988). Line drawings contain sufficient visual information to allow humans to rapidly judge perceptual and semantic aspects of scenes (Biederman, Mezzanotte, & Rabinowitz, 1982; Biederman, Teitelbaum, & Mezzanotte, 1983; Kim & Biederman, 2010; 2011). In addition to resulting in similar behavioral error patterns (Walther and Shen, 2014), color photographs and line drawings of natural scenes also elicit similar neural representations of scene categories in the PPA (Walther et al., 2011). More importantly, line drawings provide explicit descriptions of several informative contour properties not readily accessible in full-textured color photographs, such as contour orientation, length, curvature, and types and angles of junctions created by multiple contours (Walther & Shen, 2014). The current study benefits from this direct access to important contour properties by manipulating predictability of either orientation or junction statistics for scene categories.

We tested the causal role of these two sets of candidate features, orientation statistics and junction properties, for the neural representation of scene categories in the human brain. Scene categories were decoded from the fMRI activity of participants, who passively viewed blocks of line drawings of six scene categories: beaches, forests, mountains, city streets, highways, and offices. A block design was employed for its robust signal and for its proven capability to detect category-specific signals in common to multiple stimuli in a block (Cox & Savoy, 2003; Epstein & Kanwisher 1998; Haxby, Gobbini, Furey, Ishai, Schouten, & Pietrini, 2001; Kim & Biederman, 2010; Park et al., 2011; Walther et al., 2009). We devised two image manipulations that allowed for selective disruption of orientation or junction statistics: One is to rotate line drawings by random angles, which selectively disrupts orientation statistics. The other is to shift randomly contours of line drawings, which disrupts contour junctions. We then attempted to decode scene categories from the brain activity of participants while they viewed these manipulated images. Comparing the results to decoding scene categories from intact images allowed us to assess the causal involvement of the respective scene properties in the representation of scene categories. Note that these manipulations *alter* category-specific statistics of the targeted property to be spurious and uninformative. Although deletion of junctions could be a direct manipulation, pixel removal around junction locations inevitably affects non-targeted contour properties, such as statistics of orientation, curvature and length of contours. Our image manipulations, therefore, were to ensure that only one of the two candidate properties was disrupted by each of the manipulations.

We found that the category representation in two high-level visual areas involved in scene processing, the PPA, and to some extent, the occipital place area (OPA) and the lateral occipital complex (LOC) relies heavily on junction properties. It was not possible to decode scene categories from these areas when junctions were disrupted. Disrupting orientation statistics, on the other hand, did not affect the representation of scene categories. By contrast, scene categories could be decoded from neural activity patterns in early visual cortex well above chance for images with disrupted junction or orientation statistics just as from intact images. We further found that correlation of decoding error patterns from the PPA with behavioral error patterns was contingent on the preservation of junction properties, whereas disrupting orientations had no effect on error correlations. Finally, we mapped the reliance of the neural representations of scene categories on several visual properties throughout visual cortex by matching patterns of neural decoding errors to error patterns from five computational models of scene categorization from Walther and Shen (2014).

## 2. Materials and Methods

### 2.1 Participants

Sixteen healthy participants (5 females; mean age = 21.6, Standard Deviation (*SD*) = 2.8; one left-handed) were recruited from The Ohio State University community for the functional magnetic resonance imaging (fMRI) experiment, for which they received monetary compensation of $15 per hour. A separate group of 49 undergraduates at the Ohio State University participated in the behavioral experiment for course credit. Participants gave written informed consent. Both experiments were approved by the institutional review board of The Ohio State University. All participants had normal or corrected-to-normal vision and normal color vision and reported no history of neurological abnormalities. One participant (female, right-handed) was excluded from further analysis of the fMRI experiment due to excessive head movement during scans. Three participants were excluded from the analysis of the behavioral experiment because they did not complete the entire experiment or did not comply with the instructions.

### 2.2 Stimuli & Apparatus

Stimuli consisted of 475 vectorized line drawings of six categories of real-world scenes (beaches, forests, mountains, city streets, highways, and offices) from Walther et al. (2011). We generated three versions of each line drawing. Intact line drawings were generated by applying a curcular aperture (Figure 1A). Rotated line drawings were rotated the whole image by an angle randomly selected from 10 – 340° with 30° increments (Figure 1B). Distributions of contour orientation peaked at 0° and 90° in most of the six scene categories (Walther & Shen, 2014). Thus, rotation by 90°, 180° and 270° were deliberately avoided. Contour-shifted line drawings were generated by randomly translating individual contours within the circular aperture (Figure 1C). This manipulation ensured the disruption of the relations between contours, represented by contour junctions, while keeping all other contour properties constant. Note that both image rotation and random contour-shifting change local contour property statistics. Random image rotation not only systematically alters the original orientation statistics within a local image patch, but it also changes the contour junction statistics within that local patch (i.e., the contour junction statistics of a different portion of the image will substitute the original local statistics). Similarly, random contour-shifting alters orientation statistics within a local image patch, and also generates new spurious contour junctions in the patch. The difference between these manipulations is their effect on *global* image statistics. Rotation preserves the 3D relationships between surfaces and objects, and contour shifting does not.

**Figure 1.**
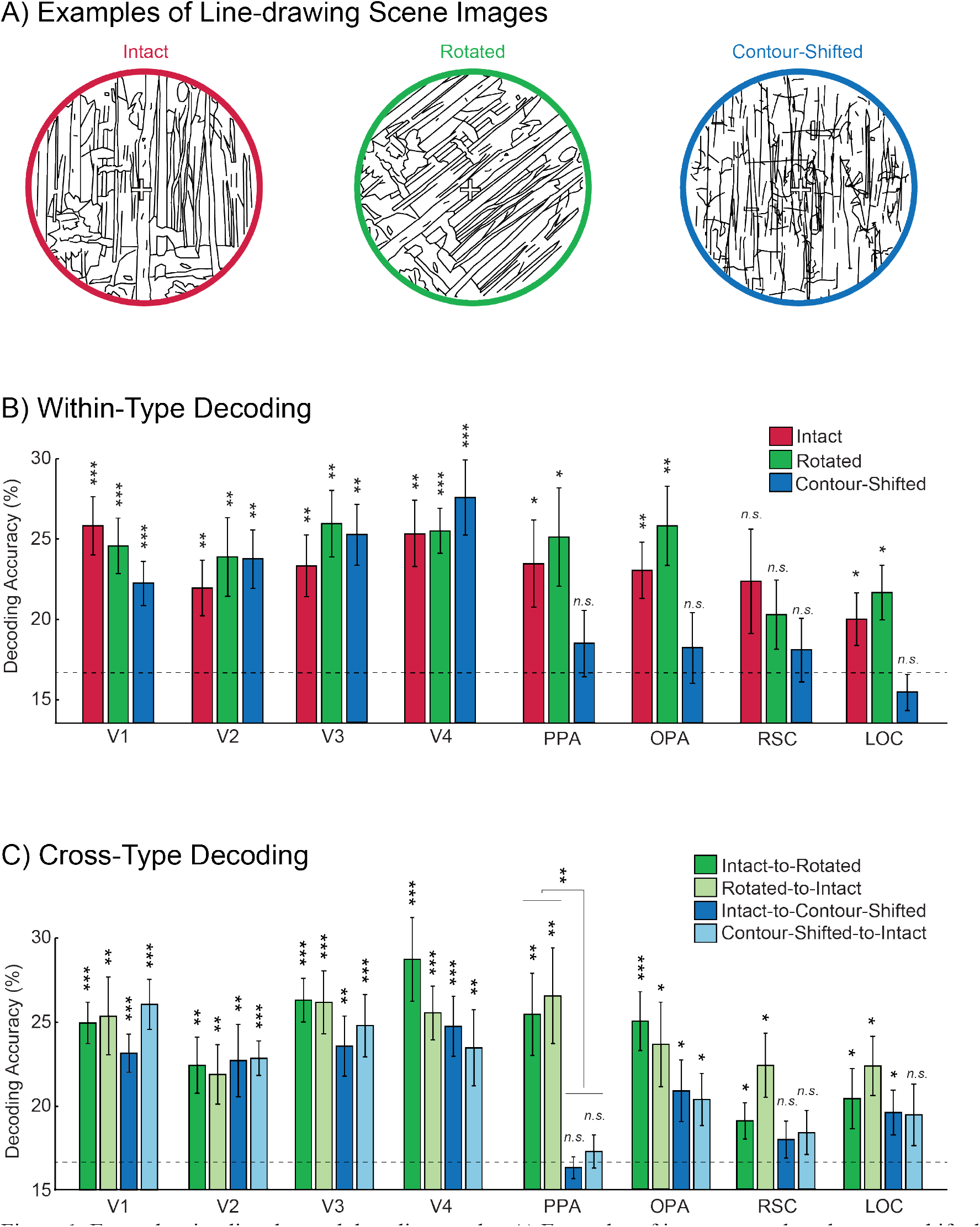
Example stimuli and neural decoding results. A) Examples of intact, rotated, and contour-shifted line drawings were derived from the same color photograph of a forest scene. The set of contours is identical across the three images within a triplet. The circular outline is shown here only for illustration and was absent in the stimuli seen by participants. B) Average accuracy rates of within-type category decoding from ROIs. C) Average accuracy rates of cross-type category decoding from the ROIs. The only significant difference in cross-type decoding accuracy between Intact-to-Rotated/Rotated-to-Intact and Intact-to-Contour-Shifted/Contour-Shifted-to-Intact was found in the PPA, as indicated above the bracket bridging the two bars. Error bars are standard errors of means. Dashed lines indicate chance performance (1/6). The significance of the one-sample t-test (one-tailed) was adjusted for multiple comparisons and marked above each bar, \**q* < .05, \*\**q* < .01, \*\*\**q* < .001.

As a result of these two manipulations, we obtained a total of 475 triplets, each of which consisted of an *intact,* a *rotated*, and a *contour-shifted* line drawing derived from the same color photograph of a real-world scene^1^. The same triplets were used for all participants in both fMRI and behavioral experiment.

Both experiments were controlled using Python 2.5 with VisionEgg 1.2 on a PC with Microsoft Windows XP. Stimuli for the fMRI experiment were back-projected onto a screen mounted in the back of the scanner bore with a DLP projector (Christie DS+6K-M 3-chip SXGA+) at a resolution of 1280 x 1024 pixels. Participants viewed stimuli through a mirror mounted on the head coil. Line drawings were rendered as black lines on a white background (2 pixels width) at a resolution of 1023 x 1023 pixels, which subtended approximately 17° x 17° of visual angle. Line drawings were seen through a circular aperture of 1023 pixels diameter. The part of the screen outside the circular aperture was 50% gray. A white fixation cross with a black outline was drawn at the center of the screen and subtended 0.5° x 0.5° of visual angle. Stimuli for the behavioral experiment were displayed on a CRT monitor with 1024 x 768 pixels resolution and 150 Hz refresh rate. Line drawings were rendered as black lines of 1-pixel width on a white background at a resolution of 600 x 600 pixels (approximately 18° x 18°), and seen through a circular aperture of 600 pixels diameter. The fixation cross had a size of 1° x 1°.

### 2.3. Experiment Design

In the fMRI experiment, participants were asked to attentively view the line drawings while fixating on the central cross. To ensure that participants followed the instruction, we monitored participants’ eye-movements in real-time using an MR compatible Eyelink 1000 system, but eye-movements were not recorded. Each participant viewed a total of 384 triplets (64 per scene category) randomly chosen from the 475 triplets. All participants had eight runs, 6 min and 12 sec in length. Each run included 18 blocks of all possible combinations of the three line drawing types (intact, rotated, and contour-shifted), and the six basic scene categories (beaches, forests, mountains, city streets, highways, and offices). During each block, eight line drawings from the same image type and scene category were shown for 800 ms, followed by a 200 ms blank per scene image. 12 sec of a blank fixation periods were inserted between blocks as well as at the beginning and the end of each run. The order of blocks within runs was counterbalanced across runs and participants according to image type and scene category.

In the behavioral experiment images from all six scene categories were randomly interleaved. Participants were asked to indicate the categories of scene images by pressing one of six keys (s, d, f, j, k, and l) on a computer keyboard. The mapping between categories and keys was assigned randomly to each participant. Each trial started with a fixation period of 500 ms, followed by a line drawing for a variable amount of time (250 ms initially), which was followed by a texture mask for 500 ms and a blank period for another 2000 ms. The texture mask was derived from a mixture of textures synthesized from all six scene categories (Loschky, Hansen, Sethi, & Pydimarri, 2010; Portilla & Simoncelli, 2000). Participants’ key responses were recorded from the onset of the image until the end of the blank period. If no response was made by the end of the blank period, the trial was recorded as incorrect. In the first phase of the experiment, participants practiced the response mapping until they achieved 90% accuracy. In the following stair-casing phase the stimulus onset asynchrony (SOA) was adjusted to 65% accuracy using the QUEST algorithm (Watson & Pelli, 1983) for each participant. By using the stair-cased SOA and the perceptual masking procedure, we aimed to provoke erroneous responses, so that we could compare the error patterns between behavior and neural decoding reliably. A randomly selected subset of twelve triplets of each category was shown for practice and stair-casing, leaving 60-68 triplets per category for testing. In practice and stair-casing, images were presented in their intact version, and participants were alerted to their mistakes by a beep.

During the testing phase, 60 triplets per category were randomly selected from the unused sets. Each image was shown only once during testing, either as an intact line drawing, a rotated line drawing, or a line drawing with randomly contour-shifted contours. 360 trials were grouped into 18 blocks of 20 images. All three line drawing types and six scene categories were presented intermixed within a block. Participants no longer received feedback during the testing phase of the experiment. The SOA was fixed to the final SOA of the stair-casing procedure. We excluded data from the seven participants with SOAs exceeding 100 ms from further analysis^2^. As a result, the final SOAs of the remaining 39 participants ranged from 17 – 87 ms (*M* = 34 ms, *SD* = 19 ms). The behavioral data were recorded in three confusion matrices, one for each image type. The rows of a confusion matrix indicate the scene categories presented to a participant, and the columns indicate the participant’s response. Cells contain the relative frequency of participants responding with the category indicated by the column, given that the presented image was of the category indicated by the row. Thus, diagonal entries contain correct responses of a scene categorization task, and off-diagonal entries contain errors in scene categorization.

### 2.4. fMRI Data Acquisition and Preprocessing

MRI images were recorded on a 3 Tesla Siemens MAGNETOM Trio MRI scanner with a 12-channel head coil at the Center for Cognitive and Behavioral Brain Imaging (CCBBI) at The Ohio State University. High-resolution anatomical images were obtained with a 3D-MPRAGE (magnetization-prepared rapid acquisition with gradient echo) sequence with sagittal slices covering the whole brain; inversion time = 930 ms, repetition time (TR) = 1900 ms, echo time (TE) = 4.44 ms, flip angle = 9°, voxel size = 1 x 1 x 1 mm, matrix size = 224 x 256 x 160 mm. Functional images were recorded with T2*-weighted echo-planar sequences with coronal slices, covering approximately the posterior 70% of the brain: for the main experiment, TR = 2000 ms, TE = 28 ms, flip angle = 72°, voxel size = 2.5 x 2.5 x 2.5 mm, matrix size = 90 x 100 x 35 mm. fMRI data were registered to a reference volume (the first volume of the fourth run) using AFNI to correct for head motion during the experiment. Then, fMRI data were smoothed using a 2 mm full-width-at-half-maximum (FWHM) Gaussian filter and converted to percentage signal change with respect to the mean of each run.

### 2.5. ROI-Based Neural Decoding

#### 2.5.1. Decoding Accuracy

As a preprocessing step for neural decoding we regressed out nuisance parameters using a general linear model (GLM) with regressors only for head motion and scanner drift.

The residuals of the GLM analysis were averaged over the durations of individual blocks, subject to a hemodynamic delay of 4 sec. The resulting 144 brain volumes (one for each block) were used as input for multi-voxel pattern analysis (MVPA).

MVPA was performed within pre-specified ROIs using a linear support vector machine (SVM) classifier (linear kernel, using LIBSVM, Chang & Lin, 2001). The classifier was trained to associate the correct category label to the blocks in seven of the eight runs, leaving out one run for testing. The categories for the blocks in the left-out run were predicted by the trained classifier. This leave-one-run-out (LORO) cross-validation was repeated until each of the eight runs was left out once. The fraction of blocks with correct test predictions was recorded as accuracy, and misclassifications were recorded in a confusion matrix (for example see Figure 2B). To investigate the effect of image rotation and contour-shifting on the representation of scene categories, LORO cross validation was performed both within image type (using the same image type for training and testing) and across image types (training on one and testing on another image type). In each case, accuracy was compared to chance (1/6) at the group level using one-tailed one-sample t-tests.

**Figure 2.**
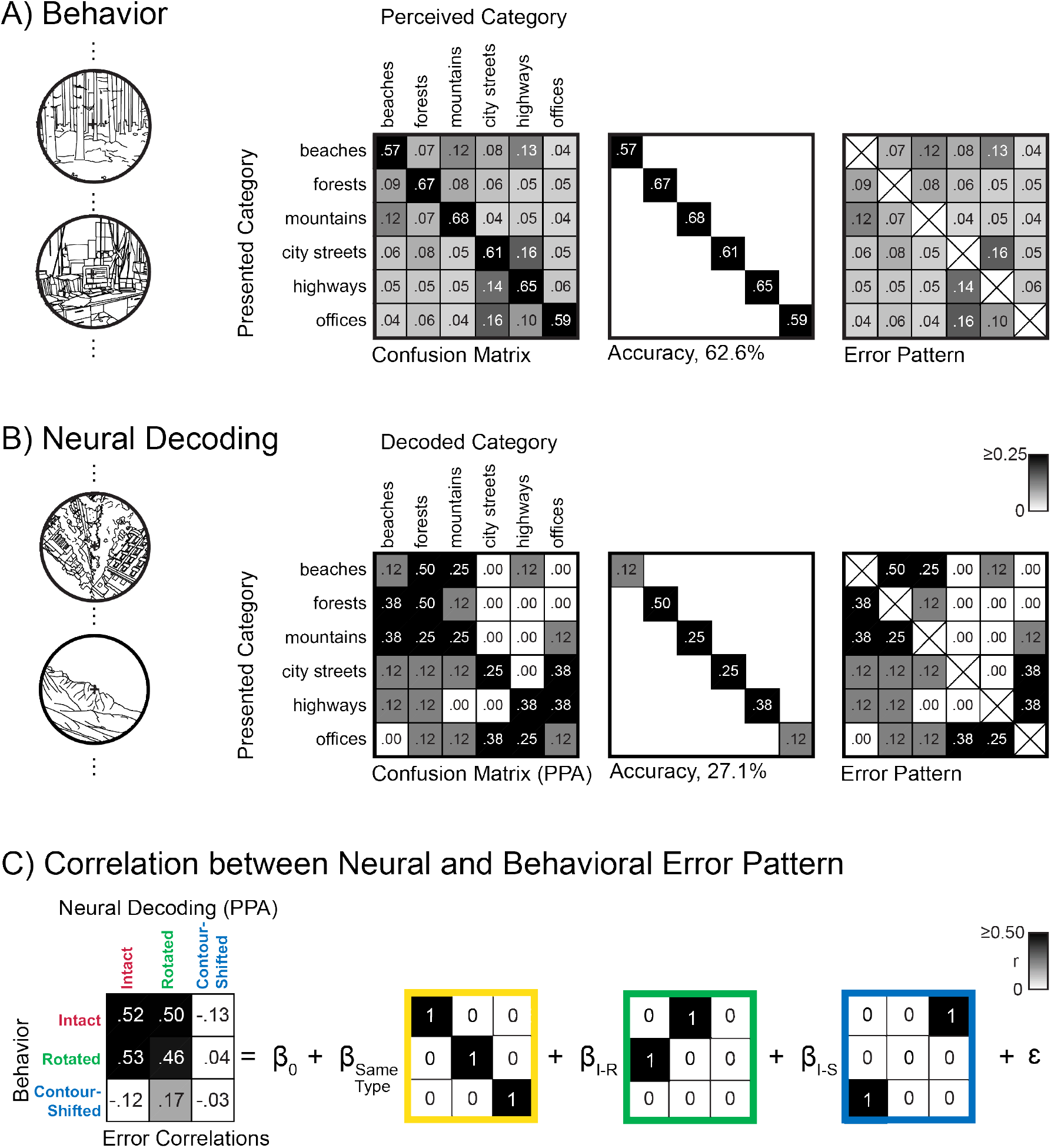
A schematic description correlation between brain and behavior error pattern. A) Group-average confusion matrix for behavioral scene categorization of rapidly presented intact line drawings. B) Confusion matrix obtained from decoding scene categories from rotated line drawings in the PPA for an individual participant. C) Off-diagonal entries of the confusion matrices were correlated for all three image types, resulting in a three-by-three error correlation matrix. The error correlations entered a linear regression analysis to measure how much each of the hypothesized models explains the observed patterns of error correlations.

#### 2.5.2. Error Pattern Correlation between Brain and Behavior

We measured the similarity of underlying categorical representations by correlating decoding error patterns from each of the ROIs to behavioral error patterns from the rapid scene categorization experiment (Walther, Beck, & Fei-Fei, 2012). The vector consisting of the 30 off-diagonal entries of the confusion matrix from an ROI was correlated to the vector of 30 off-diagonal entries of the confusion matrix from behavioral scene categorization. Statistical significance of the correlation was established non-parametrically against the null distribution of all error correlations obtained from jointly permuting rows and columns of the behavioral confusion matrix.

How does disruption of contour orientation or junction properties modulate error correlations between neural decoding and behavior? Since we had the same three image types (i.e., intact, rotated, and contour-shifted) for both neural decoding and behavioral rapid scene categorization, we could examine effects of property disruption in a 3-by-3 pattern of correlations. We modeled these correlation patterns as a linear combination of three models of the interaction between behavior and patterns of brain activity (Figure 4C). The first model is straightforward in that error correlations between neural decoding and behavior for the same image types are correlated (*same type* model). The second model states that error patterns from neural decoding for intact line drawings correlate to error patterns from the behavioral categorization of rotated line drawings and vice versa. This *intact–rotated* model prediction is based on the idea that junction properties are necessary for maintaining the error pattern similarities between neural decoding and behavior, which had been reported using intact line drawings (Walther et al. 2011). The last model assumes that orientation statistics leads to similarities in error patterns between neural decoding and behavior. This *intact–contour-shifted* model predicts high error correlation between neural decoding and behavior between intact and contour-shifted line drawings. The weights belonging to the three models were obtained by linearly regressing the error correlation patterns for all MRI participants onto the model regressors shown in Figure 2C.

**Figure 3.**
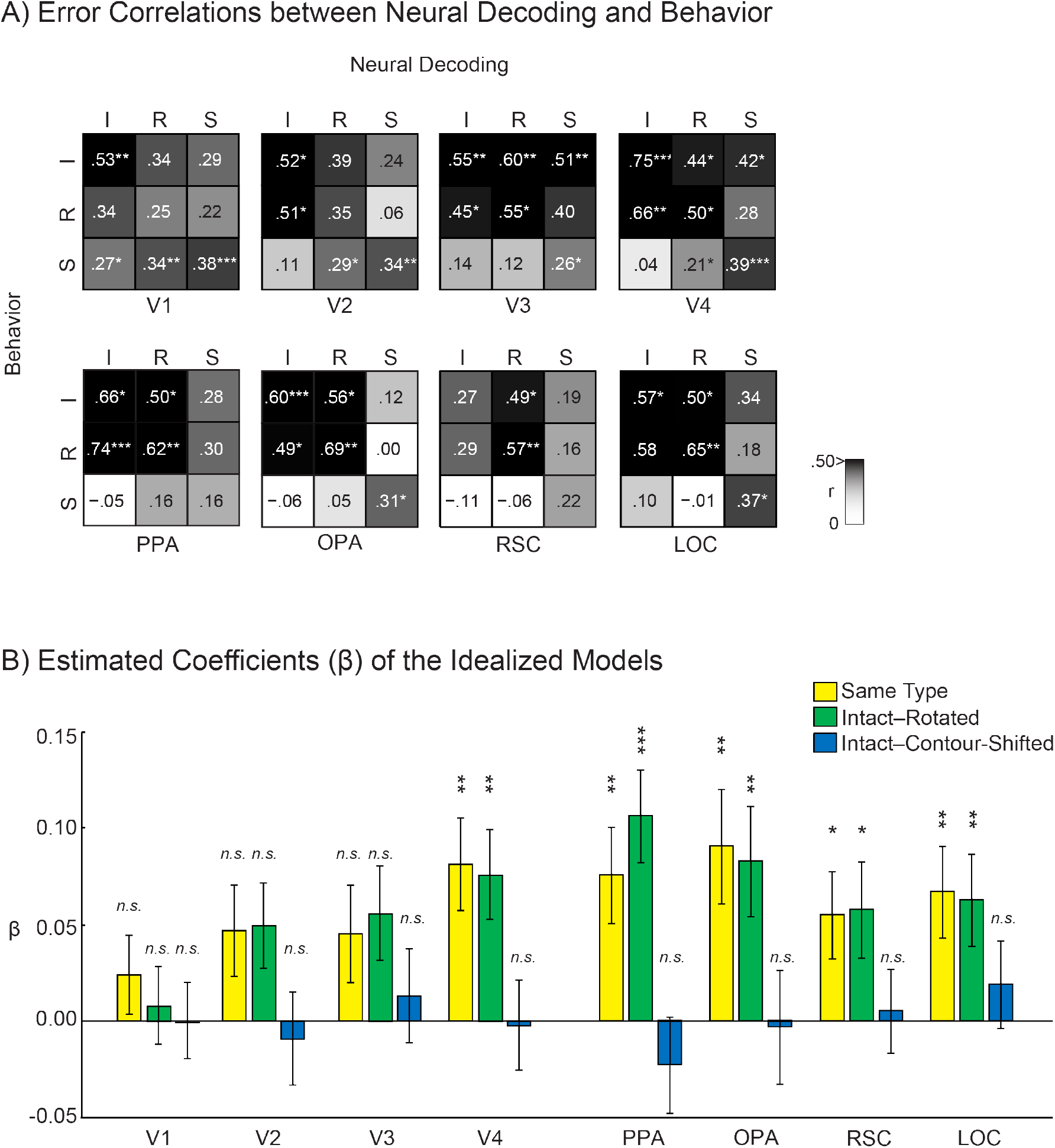
Results of error pattern correlations between neural decoding and behavior. A) Error correlations between neural decoding and behavior in V1-4, PPA, OPA, RSC, and LOC. Three-by-three correlation matrices were created by correlating group average confusion matrices obtained from fMRI and group average confusion matrices from the behavioral experiment. I stands for intact, R for rotated, and S for contour-shifted line drawing conditions for both neural decoding and behavior. The rows represent behavioral conditions, and the columns represent neural decoding conditions. Thus, each entry of a correlation matrix indicates an error pattern correlation value, r, between neural decoding and behavior. The significance of the correlation was determined non-parametrically using a permutation test, in which we computed correlations for all 720 permutations of the six category labels. B) Estimated coefficients of the three idealized models obtained from the ROIs. Error bars are estimated standard errors of means. The significance of the one-sample t-test (two-tailed) was corrected for multiple comparisons using false discovery rate and marked above each bar, \**q* < .05, \*\**q* < .01, \*\*\**q* < .001.

**Figure 4.**
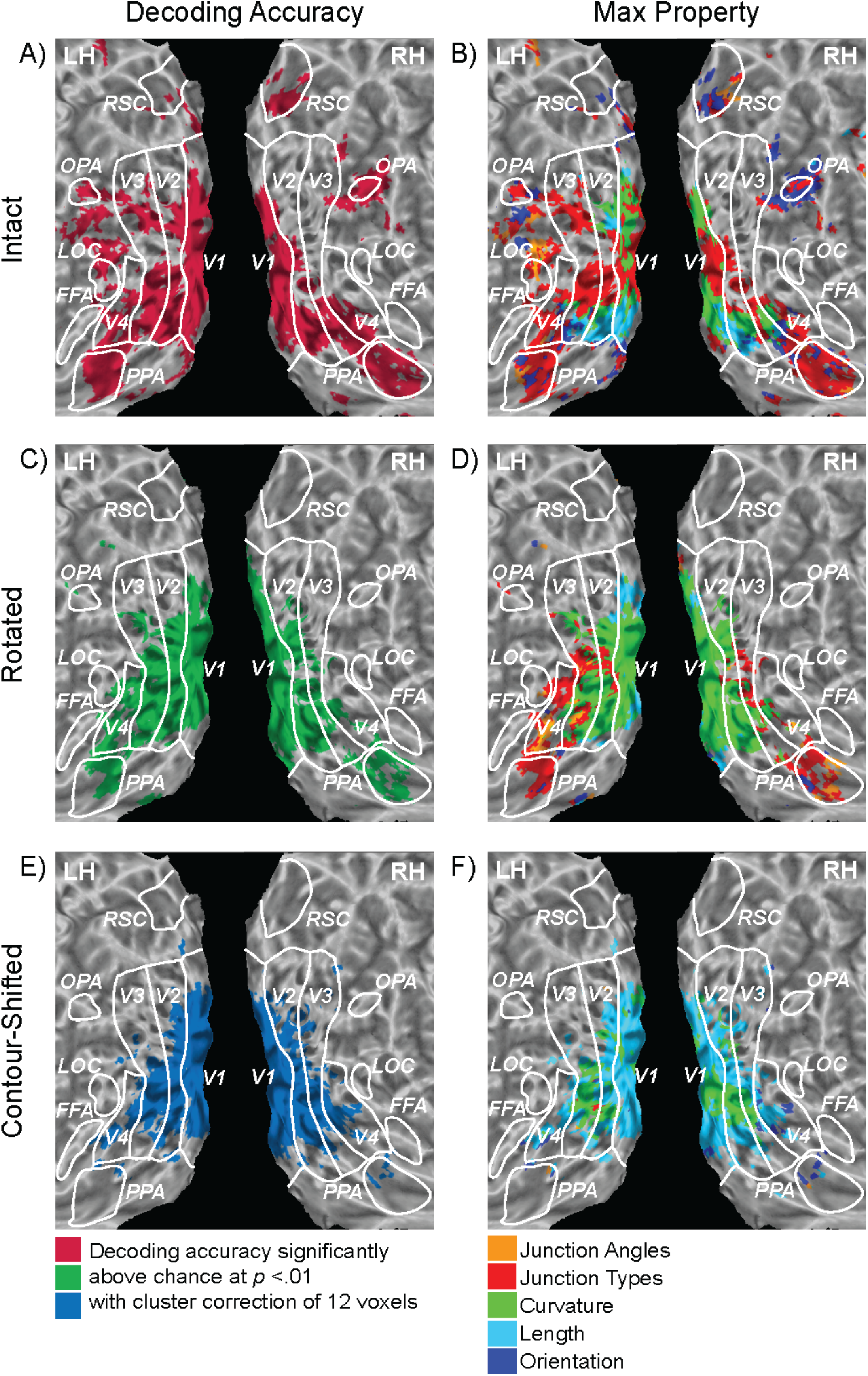
Neural decoding accuracy and max property maps. A) Searchlight locations with above-chance decoding of intact line drawings are highlighted in red. B) Property with the highest error correlation between searchlight decoding and computational feature model (max property) at the searchlight locations with above-chance decoding of intact line drawings. C) Searchlight locations with above-chance decoding of rotated line drawings are highlighted in green. D) Max property at the searchlight locations with above-chance decoding of rotated line drawings. E) Searchlight locations with above-chance decoding of contour-shifted line drawings are highlighted in blue. F) Max property at the searchlight locations with above-chance decoding of contour-shifted line drawings.

We tested which of the three models explain the relationships in the 3-by-3 error correlation patterns using a linear mixed-effects model (lme4 package in R, Bates, Maechler, Bolker, & Walker, 2014), which included the three idealized models as fixed effects, and participants as random effects. Prior to the regression analysis, error pattern correlation values were normalized using Fisher’s z transformation. Thus, the coefficients of the predictors provided estimates of how well each of the three models predicts the error correlations between neural decoding and behavior.

### 2.6. Searchlight Analysis

#### 2.6.1. Decoding Accuracy

We explored how the human visual cortex outside of the pre-defined ROIs represents categorical information about scenes using the Searchmight toolbox (Pereira & Botvinick, 2011). Searchlight analysis was performed with partial coverage in the coronal direction, which was sufficient to encompass approximately the posterior 70% of the brain on average across the participants. The same block-averaged data used in the previous ROI-based analysis entered the searchlight analysis. We defined a cubic “searchlight” of 125 voxels, whose size was matched to the average size of unilateral PPA across participants (142.9 voxels, *SD* = 66.8 voxels) as closely as possible. The searchlight was centered on each voxel at a time (Kriegeskorte, Göbel, & Bandettini, 2006), and LORO cross-validation analysis was performed within each searchlight location using a Gaussian Naïve Bayes classifier until all voxels served as the center of the searchlight. Decoding accuracy, as well as the full confusion matrix at a given searchlight location, were assigned to its central voxel. We performed the searchlight analysis separately for decoding of each of the three line drawing types, resulting in three individual accuracy maps for each participant.

To examine the agreement between the searchlight and the ROI-based analysis, we counted overlap between searchlight results and the areas V1-4, PPA, OPA, RSC, LOC, and FFA. Accuracy maps were thresholded at *p* < .005 (one-tailed p-values were obtained by the analytic methods provided by the Searchmight toolbox), and cluster-corrected using α probability simulation separately per each participant. Overlap with ROIs was computed as a percentage of ROI voxels that were included in the thresholded accuracy maps.

For group-analysis, we first co-registered each participant’s anatomical brain to the Montreal Neurological Institute (MNI) 152 template (Fonov, Evans, McKinstry, Almli, & Collins, 2011) using a diffeomorphic transformation as calculated by AFNI’s 3dQWarp. We then used the same transformation parameters to register individual decoding accuracy maps to MNI space using 3dNWarpApply, followed by spatial smoothing with a 2 mm FWHM Gaussian filter. To identify voxels with decodable categorical information, we performed one-tailed t-tests to test whether decoding accuracy at each searchlight location was above chance (1/6). After thresholding at *p* < .005 (one-tailed) we conducted a cluster-level correction for multiple comparisons, applying a minimum cluster size of 12 voxels, the average cluster size obtained from the α probability simulations conducted individually (*SD* = 0.7 voxels). After thresholding by decoding accuracy, error pattern correlations were computed between decoding from a searchlight location and each of the five computational models to determine max property of each searchlight location.

The group-level ROIs were drawn by registering ROIs of individual participants to MNI space using the same transformation parameters and overlaying them. Voxels counted in at least four participants were defined as group-level ROIs. This decision was made to ensure reasonably sized group-level ROIs while minimizing overlap between them. Finally, we excluded any voxels counted in more than a single group-level ROI.

#### 2.6.2. Error Pattern Correlation to Computational Models

We asked which contour properties contribute to the categorical representations contained in each searchlight location by comparing error patterns (Walther, Beck, & Fei-Fei, 2012). Previously, we had developed computational descriptions of the same line drawings based on five contour properties – orientation, length, curvature, junction types, and junction angles (Walther & Shen, 2014). These five image properties were computed directly from the vectorized line drawings. Separate linear support vector machine classifiers were trained to predict categories of line drawings of scenes using histograms of each of these five properties in turn. Test errors from a ten-fold cross-validation analysis were recorded in separate confusion matrices, one for each image property. Here, we correlated error patterns from decoding scene categories at each searchlight location with those from the computational analysis based on each of the five properties. We call the property with the highest correlation at a given searchlight location “max property”. This analysis was performed separately for each type of line drawing and restricted to voxels with above-chance decoding accuracy.

## 3. Results

### 3.1 ROI-Based Neural Decoding

#### 3.1.1. Within-Type Decoding

Separate classifiers were trained to discriminate scene categories based on neural activity patterns recorded while participants viewed intact, rotated, or contour-shifted line drawings. The classifiers were then tested on independent data in an LORO cross-validation procedure, separately for V1-4, PPA, OPA, RSC, LOC, and FFA. Consistent with previous findings, one-tailed t-tests showed that scene categories of intact line drawings were correctly decoded significantly above chance (1/6) in most of the visually active ROIs: V1-4, PPA, OPA, and LOC. Decoding accuracy for the RSC was comparable to the previously reported accuracy from the same 6-way category decoding (Walther et al., 2011), but failed to reach significance. As expected, decoding accuracy from the FFA was not significantly above chance.

How does disruption of orientation or junction statistics affect the neural representation of scene categories? In V1-4, scene categories could be decoded significantly above chance for both rotated and contour-shifted just as for intact line drawings. In the PPA, OPA, and the LOC, however, scene categories could only be decoded from rotated line drawings, but not from contour-shifted line drawings. The RSC showed a similar pattern of results, but category decoding for rotated line drawings was only marginally above chance (for average decoding accuracies and standard error of means see Figure 1B, for detailed statistical results see Table 1).

**Table 1.** Results of one-tailed t-tests for within-type decoding. Significance was adjusted using the false discovery rate for multiple comparisons correction.

| ROI | df | Within-Type Decoding |  |  |  |  |  |
| --- | --- | --- | --- | --- | --- | --- | --- |
|  |  | Intact |  | Rotated |  | Contour-Shifted |  |
|  |  | *t* | *q* | *t* | *q* | *t* | *q* |
| V1 | 14 | 5.047 | $2.67 \cdot 10^{-4}$ | 4.551 | $3.40 \cdot 10^{-4}$ | 4.044 | $6.04 \cdot 10^{-4}$ |
| V2 | 14 | 3.041 | $5.37 \cdot 10^{-3}$ | 2.941 | $5.37 \cdot 10^{-3}$ | 3.900 | $2.40 \cdot 10^{-3}$ |
| V3 | 14 | 3.453 | $1.94 \cdot 10^{-3}$ | 4.491 | $3.81 \cdot 10^{-3}$ | 4.544 | $3.81 \cdot 10^{-3}$ |
| V4 | 12 | 3.892 | $1.07 \cdot 10^{-3}$ | 5.825 | $1.22 \cdot 10^{-4}$ | 4.324 | $7.42 \cdot 10^{-4}$ |
| PPA | 14 | 2.492 | .0194 | 2.763 | .0194 | .871 | .199 |
| OPA | 11 | 3.276 | $5.54 \cdot 10^{-3}$ | 3.325 | $5.54 \cdot 10^{-3}$ | .635 | .269 |
| RSC | 14 | 1.746 | .0882 | 1.667 | .0882 | .698 | .248 |
| LOC | 14 | 2.037 | .0458 | 2.944 | .0160 | -1.108 | .857 |
| FFA | 14 | .575 | .431 | .000 | .500 | .636 | .431 |

#### 3.1.2. Cross-Type Decoding

If the neural representation of scene categories is indeed preserved under random image rotation but destroyed by shifting contours, then we should expect that a decoder trained on intact line drawings should be able to predict scene categories for rotated but not contour-shifted line drawings (and vice versa). We tested these predictions in several visual areas. Using the same LORO cross-validation procedure, we compared decoding performance across four conditions: intact-to-rotated (IR), rotated-to-intact (RI), intact-to-contour-shifted (IS), and contour-shifted-to-intact (SI).

Figure 1C shows the group average accuracies of the four cross-type conditions. The one-tailed t-test comparing decoding accuracy to chance showed that cross-type decoding from the early visual areas was successful for all four cross-type decoding conditions. By contrast, cross-type decoding from the PPA was significantly more accurate than expected by chance only for IR and RI, but not for IS and SI. The same pattern was also found in the RSC, although to a reduced extent. Interestingly, in the OPA, crosstype decoding was possible not only between intact and rotated line drawings but also between intact and contour-shifted line drawings, although less accurately. Similarly, accuracy from the LOC was significantly above chance for IR, RI, and IS, also marginally above chance for SI (for details on statistical tests, see Table 2).

**Table 2.** Results of one-tailed one-sample t-tests for cross-type decoding. Significance was adjusted using the false discovery rate for multiple comparisons correction.

| ROI | df | Cross-Type Decoding |  |  |  |  |  |  |  |
| --- | --- | --- | --- | --- | --- | --- | --- | --- | --- |
|  |  | Intact to Rotated |  | Rotated to Intact |  | Intact to Contour-Shifted |  | Contour-Shifted to Intact |  |
|  |  | *t* | *q* | *t* | *q* | *t* | *p* | *t* | *q* |
| V1 | 14 | 6.738 | $1.86 \cdot 10^{-5}$ | 3.757 | $1.06 \cdot 10^{-3}$ | 5.778 | $3.18 \cdot 10^{-5}$ | 6.330 | $1.86 \cdot 10^{-5}$ |
| V2 | 14 | 3.410 | $4.23 \cdot 10^{-3}$ | 2.882 | $7.60 \cdot 10^{-3}$ | 2.764 | $7.60 \cdot 10^{-3}$ | 5.955 | $7.02 \cdot 10^{-5}$ |
| V3 | 14 | 7.378 | $6.92 \cdot 10^{-6}$ | 5.013 | $1.90 \cdot 10^{-4}$ | 3.780 | $1.02 \cdot 10^{-3}$ | 4.326 | $4.64 \cdot 10^{-4}$ |
| V4 | 12 | 4.484 | $7.48 \cdot 10^{-4}$ | 5.126 | $5.02 \cdot 10^{-4}$ | 4.156 | $8.90 \cdot 10^{-4}$ | 2.756 | $8.70 \cdot 10^{-3}$ |
| PPA | 14 | 3.572 | $3.89 \cdot 10^{-3}$ | 3.452 | $3.89 \cdot 10^{-3}$ | -.642 | .734 | .564 | .388 |
| OPA | 11 | 4.231 | $2.82 \cdot 10^{-3}$ | 2.469 | .0312 | 2.031 | .0336 | 2.093 | .0336 |
| RSC | 14 | 2.200 | .0451 | 2.978 | .0120 | 1.126 | .139 | 1.261 | .139 |
| LOC | 14 | 2.099 | .0363 | 3.246 | .0112 | 2.193 | .0363 | 1.512 | .0752 |
| FFA | 14 | -.105 | .631 | -.341 | .631 | -.188 | .631 | 1.183 | .513 |

To further examine the differences across the brain regions, we conducted a repeated measures ANOVA by using two factors: (1) Which *type* of disruption was used for the cross-type decoding, rotated (IR and RI) vs. contour-shifted (IS and SI), and (2) in which *direction* the decoding was conducted, trained on intact line drawings and tested on disrupted line drawings (IR and IS) vs. trained on disrupted line drawings and tested on intact line drawings (RI and SI). Consistent with the results from the one-sample t-tests, the main effect of type was significant in the PPA, *F*(1, 14) = 17.681, *p* = 8.82·10^‐4^, *η*^*2*^ = .558. In other scene-sensitive areas, however, it failed to reach significance: in the OPA, *F*(1, 11) = 1.983, *p* = .188, *η*^*2*^ = .152, in the RSC, *F*(1, 14) = 3.278, *p* = .0917, *η*^*2*^ = .190, and in the LOC, *F* < 1. As expected, in the early visual areas the main effect of type was not significant; *Fs* < 1 in V1-3, and *F*(1, 12) = 1.370, *p* =.265, *η*^*2*^ = .102 in V4. Neither the main effect of *direction* nor the interaction between *type* and *direction* was significant in any of the ROIs.

#### 3.1.3. Correlation between Neural and Behavioral Error Pattern

To explore the similarity of the underlying categorical representations between neural decoding and behavior, we performed a behavioral categorization experiment with a separate group of 49 participants. Participants were shown a line drawing of natural scenes, followed by the perceptual mask, and asked to indicate its scene category as either a beach, a forest, a mountain, a city street, a highway, or an office. Following practice and stair-casing, participants’ performance stabilized at stimulus-onset-asynchronies (SOA) of 13 – 87 ms (*M* = 34 ms, *SD* = 19 ms). Average accuracy during the test phase pooled over all line drawing types was 45.5 % (Standard Errors of Means (*SEM*) = 1.8%). A repeated-effects ANOVA of accuracy showed a significant effect for type of line drawing, *F*(1.69, 64.06) = 148.476, *p* = 2.22·10^-16^, *η*^*2*^ = .796 (degrees of freedom were adjusted due to a violation of sphericity). Accuracy was highest for intact line drawings (*M* = 62.3%, *SEM* = 2.9%), followed by rotated line drawings (*M* = 44.1%, *SEM* = 2.2%), and lowest for contour-shifted line drawings (*M* = 30.0%, *SEM* = 1.2%). The accuracy of contour-shifted line drawings was still significantly above chance, *t*(38)=11.051, *p* = 9.88·10^‐14^.

Responses from the behavioral experiments were recorded in confusion matrices, separately for the three types of line drawings (see Figure 2A for intact; for behavioral confusion matrices of all three image types, see Figure S1). Off-diagonal elements of the confusion matrices represent categorization errors. Errors from the behavioral experiment were correlated with the errors made when decoding scene categories from brain activity for each of the three types of line drawings, separately per ROI and participant (see Figure 2B for the confusion matrix for decoding rotated line drawings from the PPA of one participant; for exclusive confusion matrix for neural decoding see Figure S2-S3). As can be seen in Figure 3A, error correlation was high when comparing the behavior and neural decoding for the same types of images (the diagonal of the three-by-three correlation matrices). In the PPA and OPA, the correlation was also high between intact and rotated, but not between intact and contour-shifted line drawings. In V1, no particular pattern of error correlations is discernible.

We modeled the error correlation patterns as a linear combination of three idealized models. The same-type model hypothesized that error patterns would match between behavior and decoding only for the same types of line drawings. The intact-rotated model predicts high brain-behavior error correlations between intact and rotated line drawings, assuming that disruption of orientation leaves error correlations largely unaffected. The intact-contour-shifted model, by contrast, predicts high brain-behavior error correlations between intact and contour-shifted line drawings, assuming that error correlations are maintained when junctions are disrupted, (Figure 2C).

The coefficients corresponding to each of the three idealized models were computed using a mixed-effects linear regression model with prediction models as fixed effects and participants as random effects. Figure 3B shows the estimated coefficients of the fixed effects for the prediction models for all ROIs. In V1-3, none of the three models significantly explained the error correlation patterns between neural decoding and behavior. However, further along in the visual processing stream, both the same type and the intact-rotated model significantly explained error correlation in V4, PPA, OPA, RSC, and LOC. The intact-contour-shifted model, on the other hand, did not contribute to the patterns in any of the ROIs (for details on statistical tests see Table 3).

**Table 3.** Results of two-tailed t-tests for coefficients of the three idealized models for explaining patterns of error correlation between neural decoding and behavior. Significance was adjusted using false discovery rate for multiple comparisons correction.

| ROI | *df* | Same Type |  | Intact-Rotated |  | Intact-Contour-Shifted |  |
| --- | --- | --- | --- | --- | --- | --- | --- |
|  |  | *t* | *q* | *t* | *q* | *t* | *q* |
| V1 | 14 | 1.166 | .732 | .391 | .999 | -1.646·10<sup>-5</sup> | .999 |
| V2 | 14 | 1.878 | .0718 | 2.229 | .0718 | -.103 | .918 |
| V3 | 14 | 1.786 | .111 | 2.289 | .0663 | -.912 | .362 |
| V4 | 12 | 3.376 | 1.10·10<sup>-3</sup> | 3.267 | 3.27·10<sup>-3</sup> | -.103 | .918 |
| PPA | 14 | 3.035 | 3.62·10<sup>-3</sup> | 4.440 | 2.69·10<sup>-5</sup> | -.912 | .362 |
| OPA | 11 | 3.056 | 5.61·10<sup>-3</sup> | 2.899 | 5.61·10<sup>-3</sup> | -.107 | .915 |
| RSC | 14 | 2.439 | .0312 | 2.311 | .0312 | .238 | .812 |
| LOC | 14 | 2.831 | 8.31·10<sup>-3</sup> | 2.774 | 8.31·10<sup>-3</sup> | .795 | .427 |
| FFA | 14 | .568 | .570 | 1.315 | .189 | -.0324 | .974 |

### 3.2. Searchlight Analysis

#### 3.2.1 Decoding Accuracy

To further characterize the representation of scene categories throughout visual cortex, we performed a searchlight analysis of the posterior 70% of the brain that was included in the partial acquisition scans. At each searchlight location, we attempted to decode scene categories using LORO cross-validation, separately for intact, rotated and contour-shifted line drawings. This analysis resulted in spatial maps of decoding accuracy and confusion matrices for each location. To assess the agreement between the searchlight analysis and the ROI-based analysis we computed the percentage of voxels in each of the ROIs that overlapped with the searchlight accuracy maps separately for each participant. The average of the amount of overlap is shown in Table 4. The searchlight maps for decoding intact line drawings showed the largest amount of overlap with all ROIs. More importantly, the overlap of searchlight maps with the PPA and the OPA was larger for rotated than for contour-shifted line drawings. By contrast, a similar amount of overlap was found for V1-4. In fact, the accuracy map of contour-shifted line drawings overlapped with slightly more V1 and V2 voxels than the accuracy map of rotated line drawings.

**Table 4.**
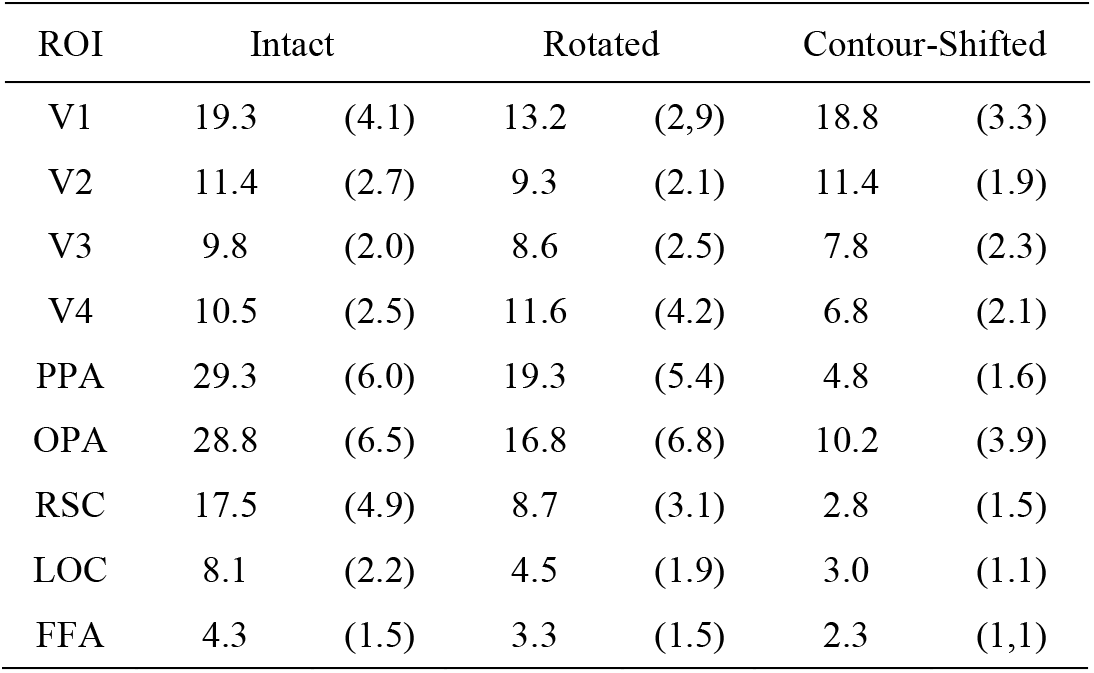
Average percentages (%) of overlap of each ROI with searchlight maps computed for the three image types. Standard errors of means are shown in parentheses.

The group-mean accuracy map of decoding intact line drawings (Figure 4A) showed a large cluster of voxels in the posterior visual cortex, including the bilateral parahippocampi, left precuneus, and the lateral end of the left transverse occipital sulcus, as well as the bilateral fusiform and calcarine gyri. This large cluster extended to the cerebellum bilaterally, but more to the left than the right cerebellum. The second large cluster encompassed the right middle occipital gyrus extending to the right transverse occipital sulcus. Another cluster included the right precuneus and the right posterior cingulate gyrus, which partially overlapped with the right retrosplenial cortex. The group-mean accuracy map of decoding rotated line drawings (Figure 4C) showed two clusters that largely overlapped with the accuracy map of decoding intact line drawings. One large cluster encompassed the left parahippocampal gyrus, the bilateral fusiform gyri, and bilateral calcarine gyri and extended to the bilateral cerebellum. The other cluster included the right parahippocampal gyrus. By contrast, the group-mean accuracy map of decoding contour-shifted line drawings (Figure 4E) revealed only one large cluster, which included bilateral calcarine gyri, fusiform gyri, lingual gyri, and the cuneus, and extended bilaterally to the cerebellum (for an exhaustive list of peak coordinates, see Table 5).

**Table 5.** Clusters identified in the searchlight analysis for within-type decoding of intact, rotated, and contour-shifted line drawings.

| Decoding Condition | Peak (MNI coordinates) | | | Accuracy (%) | Volume (μl) | Description |
| --- | --- | --- | --- | --- | --- | --- |
|  | x | y | z |  |  |  |
| Intact | 2.5 | 90.5 | -4.2 | 30.4 | 63234 | Occipital pole, calcarine gyri, fusiform gyri, parahippocampal gyri, left precuneus, left transvers occipital sulcus, bilateral cerebellum |
|  | -35.0 | 85.5 | 25.8 | 25.4 | 5219 | Right middle occipital gyrus, right transvers occipital sulcus |
|  | -25.0 | 63.0 | 18.2 | 25.6 | 1922 | Right parieto-occipital sulcus, right retrosplenial cortex |
|  | 30.0 | 45.5 | 68.2 | 22.0 | 391 | Left superior parietal gyrus |
|  | -32.5 | 30.5 | -26.8 | 19.6 | 188 | Right lateral occipito-temporal gyrus (fusiform gyrus) |
| Rotated | 10.0 | 98.0 | -14.2 | 28.0 | 50938 | Occipital pole, calcarine gyri, fusiform gyri, left parahippocampal gyrus, bilateral cerebellum |
|  | -22.5 | 50.5 | -9.2 | 26.9 | 4531 | Right parahippocampal gyrus |
|  | 27.5 | 85.5 | 33.2 | 22.7 | 375 | Left inferior parietal angular gyrus, left middle occipital gyrus |
|  | -47.5 | 80.5 | -1.8 | 21.8 | 188 | Right middle occipital gyrus |
| Contour-shifted | -15.0 | 90.5 | -6.8 | 29.3 | 54938 | Occipital pole, calcarine gyri, precuneuses, fusiform gyri, lingual gyri, bilateral cerebellum |
|  | -30.0 | 30.5 | 56.2 | 22.5 | 703 | Right precentral gyrus |
|  | 42.5 | 33.0 | 28.2 | 23.7 | 563 | Left posterior lateral fissure, left inferior parieto-supramarginal gyrus. |
|  | -27.5 | 53.0 | -9.2 | 21.9 | 484 | Right medial occipito-temporal sulcus, right lingual gyrus |
|  | 50.0 | 25.5 | 48.2 | 22.6 | 453 | Left inferior parieto-supramarginal gyrus |
|  | -2.5 | 20.5 | 76.2 | 21.7 | 422 | Right paracentral gyrus, right paracentral sulcus |
|  | -57.5 | 20.5 | 78.2 | 17.8 | 344 | Right inferior parieto-supramarginal gyrus |
|  | 10.0 | 38.0 | 55.8 | 21.7 | 250 | Left paracentral gyrus, left paracentral sulcus |
|  | -10.0 | 43.0 | 78.2 | 21.4 | 234 | Right paracentral gyrus, right paracentral sulcus |
|  | 35.0 | 93.0 | 0.8 | 21.9 | 219 | Left middle occipital gyrus |
|  | -25.0 | 88.0 | 25.8 | 21.0 | 203 | Right superior occipital gyrus. |

#### 3.2.2. Important Image Property for Neural Decoding

To determine the influence of several kinds of structural properties of line drawings on the neural representation of scene categories, we correlated error patterns from each searchlight location (that is, the off-diagonal elements of the confusion matrices) to those from five computational models of scene categorization. Each of these computational models relies exclusively on one of five structural properties: contour orientation, length, curvature, and types and angles of contour junctions (Walther & Shen, 2014). Each searchlight location was labeled according to the contour property with the highest error correlation, the *max property.* Figure 4B, D, and F show max property maps for the three image types, restricted to the locations that allowed for decoding of scene categories significantly above chance (see Figures S4-S6 for unrestricted maps of error correlations and of max property).

Representation of scene categories in early visual areas relies most heavily on contour length and curvature for intact line drawings. Junction types are particularly important in foveal regions of early visual cortex, and also the PPA, the OPA, and the RSC. For rotated line drawings, curvature dominates early visual areas, while high-level visual areas continue to rely on junction properties. For contour-shifted line drawings, early visual areas rely most on contour length and curvature. Only a few searchlight locations in high-level visual areas allow for decoding of scene categories from contour-shifted line drawings. These effects are quantified more precisely by assessing overlap of these maps with ROIs (Figure 5).

**Figure 5.**
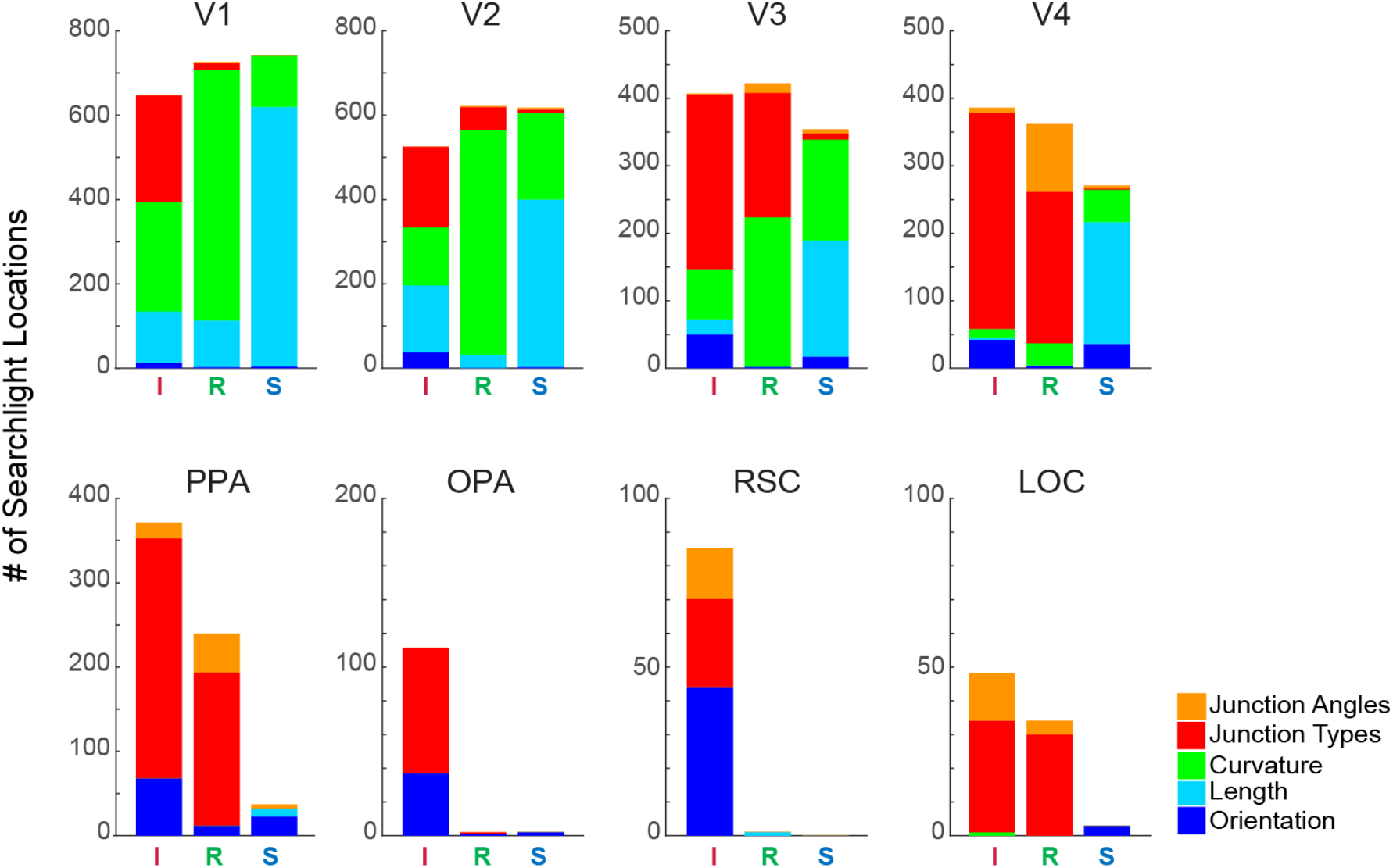
Distribution of max properties within group-level ROIs for decoding of intact (I), rotated (R), and contour-shifted (S) line drawings. Only searchlight locations with above-chance decoding accuracy were counted. Coloring follows the same convention as Figure 4.

Voxel statistics for the pre-defined ROIs show several interesting effects. First, representation of natural scene categories in V1 and V2 relies most heavily on contour length and curvature for intact line drawings. The importance of junctions increases steadily through V3 and V4, until junctions dominate the representation of scene categories in the PPA (81.6% of PPA voxels have junction angle or type as max property), the OPA (66.7%), and the LOC (97.9%). The high reliance on junction properties strongly manifested even though L-junctions, critical to surface analysis (Biederman, 1987; Guzman, 1968), were not considered in the computational category prediction (Walther & Shen, 2014). Orientation statistics, by comparison, play a minor role in the high-level visual areas, except for the RSC (51.8%).

Second, the involvement of orientation statistics in the representation of scene categories is absent for randomly rotated line drawings as expected. Whereas contour orientation was the max property for 291 out of 2575 (11.3%) searchlight locations with significant decoding for intact line drawings, this number fell to 22 out of 2402 (0.9%) searchlight locations with significant decoding for rotated line drawings. Similarly, junction properties ceased to be the max property almost everywhere for contour-shifted line drawings. Junction types and angles were the max property for 1469 out of 2575 (57.0%) searchlight locations for intact but only 38 out of 2021 (1.9%) searchlight locations for contour-shifted line drawings. These findings confirm the effectiveness of our image manipulations as a tool for probing the role of structural scene properties in the neural representation of scene categories.

Moreover, this shift in the reliance to other structural properties when deprived of contour orientation or junctions demonstrates the flexibility of early visual areas to make use of any visual regularities present in the images. However, junction properties appear to be critical for the neural representation of scene categories in high-level visual areas. Rather than shifting to other structural properties, these areas show a dramatic decrease in the number of searchlight locations that allowed for the decoding of scene categories when junctions were disrupted.

## 4. Discussion

In this study, we used multi-voxel pattern analysis (MVPA) to identify visual properties critical to the categorical representation of real-world environments in the brain. Consistent with our previous findings (Walther et al., 2009; 2011), we showed that categorical representations of scenes are distributed across the human visual cortex. Importantly, they rely on different sets of contour properties along the course of visual processing. In the PPA, OPA, and LOC, structural relations between contours as embodied by the distribution of junctions need to be preserved to maintain category-specific brain activity patterns. Whereas random shifting of contours led to chance-level decoding accuracy, random image rotation did not undermine decoding accuracy in the PPA, OPA, and LOC.

Cross-type decoding was successful between intact and rotated, but not between intact and contour-shifted line drawings in the PPA. Note that contour-shifted line drawings still preserved the statistics of contour orientation, length, and curvature. Yet, this information was not sufficient to give rise to a decodable neural representation of scene categories in the PPA, underscoring the necessity of junction properties and, thereby, structural relations between contours for the PPA to encode scene categories.

In contrast, category decoding was successful with contour-shifted line drawings as well as rotated line drawings from V1 to V4, suggesting that none of the image manipulations was detrimental to the category-specificity of neural activity patterns for line drawings in early visual areas. Furthermore, cross-type decoding was successful not only between intact and rotated but also between intact and contour-shifted line drawings, indicating that the neural representations of scene categories were compatible across the three line drawing types. In the early visual areas, category-specific neural activity patterns are not undermined by disrupting a single type of visual statistics, suggesting that the early visual areas represent scene categories only implicitly by relying on any available statistics indicative of scene categories in a parsimonious manner.

Analysis of decoding errors confirmed previous findings that error patterns of decoding from the neural activity in the PPA significantly correlate with those of human scene categorization of intact line drawings (Walther et al. 2011). Critically, we here show that junction properties but not orientation statistics of line drawings have to be preserved to maintain such brain-behavior correlation (for searchlight analysis of error pattern correlation to behavior and its results, see supplementary material; Figure S7). Preservation of junctions starts to be important for brain-behavior error correlation as early as in V2. This finding agrees well with neurophysiological studies in non-human primates showing that area V2 is sensitive to changes in types or angles of contour junctions (Pasupathy & Connor, 2002; Peterhans & von der Heydt, 1989).

In a searchlight analysis, we found early visual areas to rely on the distributions of orientation, length and curvature of contours for the neural representation of scene categories. Junction properties, on the other hand, became increasingly important in near-foveal regions of V1-4 and peaked in the PPA and OPA. The importance of junctions for the neural representation of scene categories in the PPA persisted when scene images were rotated by a random angle but disappeared when junctions were disrupted by randomly shifting contours.

As should be expected, the importance of orientation statistics for the representation of intact line drawings throughout visual cortex disappeared for rotated images and was supplanted by an increased reliance on contour length and curvature. Searchlight analysis confirms the hierarchical aspect of neural representations of scene categories. Once an image of real-world environments proceeds through the cortical hierarchy, its neural representation relies progressively more on complex visual properties, from orientation and length extracted in V1 (Hubel & Wiesel, 1962) to curvature and junction properties extracted in V2 and V4 (Pasupathy & Connor, 2002; Peterhans & von der Heydt, 1989). Recent computational work showed that junction-like feature representations arise naturally when representations of complex scenes are learned in simulated multi-layer neural networks (Zeiler & Fergus, 2014).

What would be the mechanisms for contour junctions affecting a neural representation of scene categories in the PPA? While surface features such as color and orientation gradient may initiate contour detection and surface delineation (by the primal sketch; Marr, 1982), they do not directly give a rise to the high-level representation for visual recognition. In fact, visual recognition and categorization hardly benefit from surface features once important contours are analyzed and their relations are determined (Biederman & Ju, 1988). Instead, junctions present in two-dimensional (2D) images are informative of three-dimensional (3D) structural information, because their types and angles are indicative of arrangements and relations of surfaces in 3D space (Biederman, 1987; Guzman, 1968). The 3D structure of natural environments can be inferred from the distribution of these viewpoint-invariant properties, namely junction types and angles. Furthermore, their invariance to changes in viewpoint makes these properties particularly useful for visual recognition and categorization

In fact, 3D structure of scenes is likely to be related to some global scene properties, such as whether a scene has an open or closed layout (Harel et al., 2013; Park et al., 2011). For example, scenes with an open layout are usually clutter-free and contain surfaces not obstructing one another, resulting in a relatively small number of junctions. In contrast, a closed layout is likely to contain more objects and multiple surfaces overlaying one another, thus creating proportionally more junctions. Consistent with this idea, recent neuroimaging evidence showed that the PPA is not only sensitive to changes in the statistics of simple shapes (Cant & Xu, 2012), but also to junction angles (Nasr, Echavarria, & Tootell, 2014). When combining these results with the sensitivity of the PPA to global scene properties (Kravitz et al. 2011; Park et al. 2011; Harel et al. 2013), a picture of the PPA arises as a visual area sensitive to several high-level aspects of scenes, such as semantic category, global layout, or relation of the scene to realworld locations (Marchette, Vass, Ryan, & Epstein, 2014). A recent study clearly shows that the PPA indeed can encode several aspects of a scene image such as its entry-level category, spatial layout (open vs. closed), surface texture property, and content (man-made vs. natural) (Lowe, Gallivan, Ferber, & Cant, in press). With scenes being spatial arrangements of surfaces and objects, the image properties driving these diverse visual aspects of scenes are those encoding the relationship between surfaces in 3D space, namely junction types and their angles.

Similar to the PPA, the OPA also relies more heavily on junction properties than contour orientation. Considering its anatomical proximity to the early visual cortex, the OPA is likely to subserve relatively primitive scene analysis. In fact, receptive fields in the OPA were reported to be smaller than those in the PPA (MacEvoy & Epstein, 2007). Furthermore, the OPA preferentially activates to spatial layouts over collections of multiple objects (Bettencourt & Xu, 2013; MacEvoy & Epstein, 2007). The OPA has been suggested to contain precursory representations of spatial layout of scenes based on relatively simple features (Baldassano et al., 2013; Dilks, Julian, Paunov, & Kanwisher, 2013; MacEvoy & Epstein, 2007). We here propose that contour junctions are critical for these representations.

Objects are often diagnostic for scene category (Bar & Aminoff, 2003; Greene, 2013). For instance, a beach scene is more likely to contain palm trees, beach balls, and umbrellas than desks, swivel chairs, and computer monitors, and *vice versa* in the case of an office scene. It is, therefore, natural that the object-sensitive LOC contains information about objects, which can be exploited for decoding scene category or identity (Harel et al., 2013; MacEvoy & Epstein, 2011). The LOC not only activates linearly to the number of displayed objects (MacEvoy & Epsten, 2011), but it also encodes inter-object relationships (Kim & Biederman, 2010; 2011). Given recent evidence for a strong functional relationship between the LOC and PPA, it is highly likely that information about object relations in the LOC is projected to the PPA (Baldassano et al., 2013), thus contributing the category-specific activity in the PPA. Our finding that the neural representation of scene categories in the LOC relies almost exclusively on junction properties is consistent with the reliance of invariant object recognition on non-accidental properties, most specifically contour junctions (Biederman 1987), which provide cues to the threedimensional structure of objects. In addition to their role in defining objects, contour junctions are crucial to reconstruct spatial arrangements between large-scale surfaces that define the terrain and layout of a scene, thereby contributing to the neural representations of real-world scene categories.

Random contour shifting also affected grouping properties that were not modeled explicitly in our computation analysis (Walther & Shen, 2014): proximity between parallel contours, colinearity and curvilinearity of contours indicating parallel surfaces in depth, and to some extent symmetry involving multiple contours (Biederman, 1987). These non-accidental properties have in common that they are defined by spatial relations between contours rather than by properties of individual contours. Junction statistics capture these spatial relations to only a limited extent. It is therefore remarkable that decoding error patterns in the PPA, OPA and LOC are predicted so well by junction statistics.

Unlike the other scene-selective visual regions, the RSC has been found to be critical for embedding scenes in their real-world context or memory representations rather than for perceptual analysis of scenes. For instance, the RSC plays a critical role in navigation and route learning (Aguirre & D'Esposito, 1999; Maguire, 2001), and shows preferential neural activity for landmark buildings compared to non-landmark buildings (Schinazi & Epstein, 2010). The RSC also mediates between individual scenes in a broad view rather than representing exact perceptual instances of scenes (Epstein, Parker, & Feiler, 2007; Epstein, Higgins, Jablonksi, & Feiler, 2007; Park & Chun, 2009). We found neither significant decoding accuracy nor even significant activation in the RSC for line drawings of scenes (Table S1; for detailed analysis and results see Supplementary materials). We surmise that this may be because line drawings too dissimilar from actual real-world settings that afford navigation or context-based memory retrieval.

## 5. Conclusion

We have shown that the neural representations of scene categories rely on different image properties throughout the processing hierarchy in the human visual cortex. In early visual areas, any statistical regularities available in an image had the potential to elicit category-specific patterns of neural activity. In the scene-selective high-level visual regions, especially in the PPA, accurate statistics of junction properties was necessary to generate category-specific activity patterns and, importantly, to establish high correlation of decoding error patterns with patterns of errors observed in human scene categorization behavior. We conclude that non-accidental 2D cues to 3D structure, in particular contour junctions, are causally involved in eliciting a neural representation of scene categories in the PPA and the OPA by providing a reliable description of the 3D structure of real-world environments. Summary statistics of orientations, on the other hand, are insufficient to elicit a decodable representation of scene categories in these brain regions.

## Acknowledgments

The authors declare no competing financial interests. All experiments were performed while both authors were at The Ohio State University.

## Supplementary Materials

### Localizing regions of interest

Following the main experiment scans, participants viewed blocks of color photographs of faces, scenes, objects, and grid-scrambled objects as part of a standard face-place-object localizer scan (Epstein & Kanwisher, 1998). Participants were asked to indicate immediate repetitions of images to maintain their attention to images. Images subtended 640 x 640 pixels (approximately 11 ° of visual angle) and were presented for 500 ms, followed by a blank screen for 500 ms. Participants saw five blocks of 72 images, sub-divided into four mini-blocks of 18 randomly drawn images for each of the four categories. Including 12 second fixation periods before, between, and after the blocks, the entire scan lasted for 7 minutes and 12 seconds. Scanning parameters for the face-place-object localizer differed from those of the main experiment: TR = 3000 ms, TE = 28ms, flip angle = 80°, voxel size = 2.5 x.2.5 x 2.5 mm, matrix size = 90 x 100 x 40 voxels, 40 coronal slices.

The fMRI data were motion corrected, registered to the anatomical scans that had been aligned to the functional volumes of the main experiment, spatially smoothed using a 4 mm FWHM Gaussian filter and converted to percent signal change with respect to the mean of each run. The pre-processed data entered a GLM analysis with regressors for all four image types. ROIs were defined as contiguous clusters of voxels with significant contrasts (*q* < 0.05; corrected using false discovery rate) of scenes > (faces and objects) for PPA, RSC (Epstein & Kanwisher, 1998) and OPA (Dilks et al., 2013); faces > (scenes and objects) for FFA (Kanwisher et al. 1997); and objects > (scrambled objects) for LOC (Grill-Spector, Knouf, & Kanwisher, 2004; Grill-Spector, Kourtzi, & Kanwisher 2001). To obtain robust clusters for the RSC, the threshold had to be relaxed to *p* < 0.01 (uncorrected). We could not find significant clusters corresponding to the OPA in three participants and used data of the remaining twelve participants to perform decoding from the OPA.

Boundaries of early visual areas were established by stimulating the horizontal and vertical meridians of the visual field (HV) in alternation (Kastner, Weerd, Desimone, & Ungerleider, 1998). During two HV scans (only one HV scan for four participants, because we ran out of time), participants viewed flickering checkerboard patterns (2 Hz, a mix of white, red, green, blue, and yellow checkerboards), filling pairs of wedges (width: 10°) aligned with the horizontal or vertical meridians, respectively. Fixation periods of 20 seconds were included between each alternation as well as at the beginning and the end of the scan (total scan duration: 3 min 20 sec). In order to establish the anterior boundary of V4, a scan with alternating stimulation of the upper and lower visual field (UL) with similar flickering checkerboard patterns along the diagonals was included as well, which lasted approximately 3 min. Scan parameters for the HV and UL scans were as follows; TR = 2000 ms, TE = 28 ms, flip angle = 72°, voxel size = 1.953 x 1.953 x 2 mm, matrix size = 114 x 128 x 30 voxels, 30 coronal slices.

Data from the HV and UL scans were motion-corrected, registered to the anatomical scan that had been aligned to the functional volume of the main experiment, spatially smoothed (4 mm FWHM) and converted to percent signal change. In separate GLM analyses, data from the HV scans were analyzed for a horizontal-versus-vertical meridian contrast and data from the UL scans for an upper-versus-lower visual field contrast.

Cortical surfaces for each participant’s brain were reconstructed from their anatomical scans (MPRAGE) using Freesurfer. To flatten the cortical surface, each hemisphere was virtually cut along the calcarine fissure and four additional relaxation cuts. The corpus callosum and mid-brain structures on the medial surface were removed. Boundaries between early visual areas were identified by projecting the beta-weight maps of the HV and UL contrasts onto the flattened cortical surfaces using AFNI and SUMA. Following Hansen, Kay, and Gallant (2007) we identified the V1/V2 border as the first vertical meridian, the V2/V3 border as the second horizontal meridian, and V3/V4 border as the second vertical meridian. Since V4 represents the entire contralateral hemifield on the lower bank of the calcarine fissure, we identified the anterior border of V4 as the closest boundary that encompassed both upper and lower visual field. For two participants, the anterior V4 border could not be clearly delineated. Thus, we only used data from the remaining thirteen participants to perform decoding from V4. ROIs were drawn conservatively to minimize the amount of overlap between neighboring areas. Following projection of surface-based ROIs back into the brain volume of each participant, we excluded voxels that were assigned to more than one ROI.

### Univariate analysis

To explore the effect of the disruption of contour properties on the magnitude of the average neural activity we conducted a standard general linear model (GLM) analysis using the AFNI software package, and deconvolved block responses to the three image conditions for each voxel within the ROIs. Nuisance variables were included to capture variance due to head motion (six affine transform parameters) and scanner drift (4th-degree polynomial). Beta parameters were extracted for the three contrasts: Intact > Fixation, Rotated > Fixation, and Contour-shifted >

Fixation. The beta parameters were averaged over voxels within each ROI. The GLM analysis showed that the line drawing images significantly activated all of the ROIs except for the RSC (Table S1). A one-way ANOVA showed no significant difference in neural activity between intact, rotated, and contour-shifted line drawings in any of the ROIs; *F*_*s*_ < 1 in V1, V2, PPA, RSC, and OPA; *F*(2, 28) = 1.327, *p* = .281, *η*^*2*^ = .087 in the V3; *F*(2, 24) = 1.785, *p* = .189, *η*^*2*^ = .110 in V4; *F*(2, 28) = 1.923, *p* = .165, *η*^*2*^ = .121 in the LOC; and *F*(2, 28) = 1.727, *p* = .196, *η*^*2*^= .110 in the FFA. The mean activity might not be sensitive enough to differentiate between the three types of line drawings despite some variety in neural tuning properties in the visual cortex. The line drawing scenes had naturalistic visual statistics that often contain various visual features and objects, thus equating the mean activity to an image.

**Table S1.** Results of univariate analysis. One-sample t-tests were conducted to compare average neural activity during presentation of the intact, rotated, and contour-shifted line drawings to average neural activity during fixation presentation. The mean (*M*) and standard errors of means (*SEM*), t-statistics, and the adjusted significance q are shown separately for each of the three contrasts: Intact > Fixation, Rotated > Fixation, and Contour-Shifted > Fixation.

| ROI | df | Intact > Fixation |  |  |  | Rotated > Fixation |  |  |  | Contour-Shifted > Fixation |  |  |  |
| --- | --- | --- | --- | --- | --- | --- | --- | --- | --- | --- | --- | --- | --- |
|  |  | *M* | *SEM* | *t* | *q* | *M* | *SEM* | *t* | *q* | *M* | *SEM* | *t* | *q* |
| V1 | 14 | .901 | .152 | 5.923 | $5.51 \cdot 10^{-5}$ | .872 | .146 | 5.972 | $5.51 \cdot 10^{-5}$ | .858 | .158 | 5.442 | $8.67 \cdot 10^{-5}$ |
| V2 | 14 | .855 | .140 | 6.108 | $5.66 \cdot 10^{-5}$ | .828 | .146 | 5.683 | $5.66 \cdot 10^{-5}$ | .847 | .148 | 5.725 | $5.66 \cdot 10^{-5}$ |
| V3 | 14 | .823 | .104 | 7.918 | $1.85 \cdot 10^{-6}$ | .782 | .100 | 7.793 | $1.85 \cdot 10^{-6}$ | .814 | .102 | 8.019 | $1.85 \cdot 10^{-6}$ |
| V4 | 12 | 1.15 | .127 | 8.391 | $6.90 \cdot 10^{-6}$ | 1.07 | .136 | 7.329 | $9.11 \cdot 10^{-6}$ | 1.11 | .133 | 7.770 | $7.59 \cdot 10^{-6}$ |
| PPA | 14 | .234 | .0629 | 3.723 | $5.18 \cdot 10^{-3}$ | .213 | .0645 | 3.308 | $5.18 \cdot 10^{-3}$ | .228 | .0654 | 3.486 | $5.18 \cdot 10^{-3}$ |
| OPA | 11 | .577 | .126 | 4.105 | $1.75 \cdot 10^{-3}$ | .554 | .116 | 4.282 | $1.75 \cdot 10^{-3}$ | .572 | .121 | 4.247 | $1.75 \cdot 10^{-3}$ |
| RSC | 14 | .0225 | .134 | .427 | .938 | .0107 | .135 | .242 | .938 | .0718 | .0966 | .762 | .938 |
| LOC | 14 | .572 | .113 | 5.082 | $3.86 \cdot 10^{-4}$ | .496 | .107 | 4.634 | $3.86 \cdot 10^{-4}$ | .543 | .117 | 4.640 | $3.86 \cdot 10^{-4}$ |
| FFA | 14 | .371 | .0907 | 4.088 | $1.66 \cdot 10^{-3}$ | .308 | .0747 | 4.127 | $1.66 \cdot 10^{-3}$ | .371 | .0998 | 3.716 | $2.30 \cdot 10^{-3}$ |

Moreover, the overall neural activity for any specific line drawings were likely to be smoothed out over the course of a block. These findings are consistent to the previous report that the multivariate approach can often recover the visual content despite the equivalent univariate results (e.g., Kamitami & Tong, 2005).

### Group-averaged confusion matrices of behavioral categorization and neural decoding

**Figure S1.**
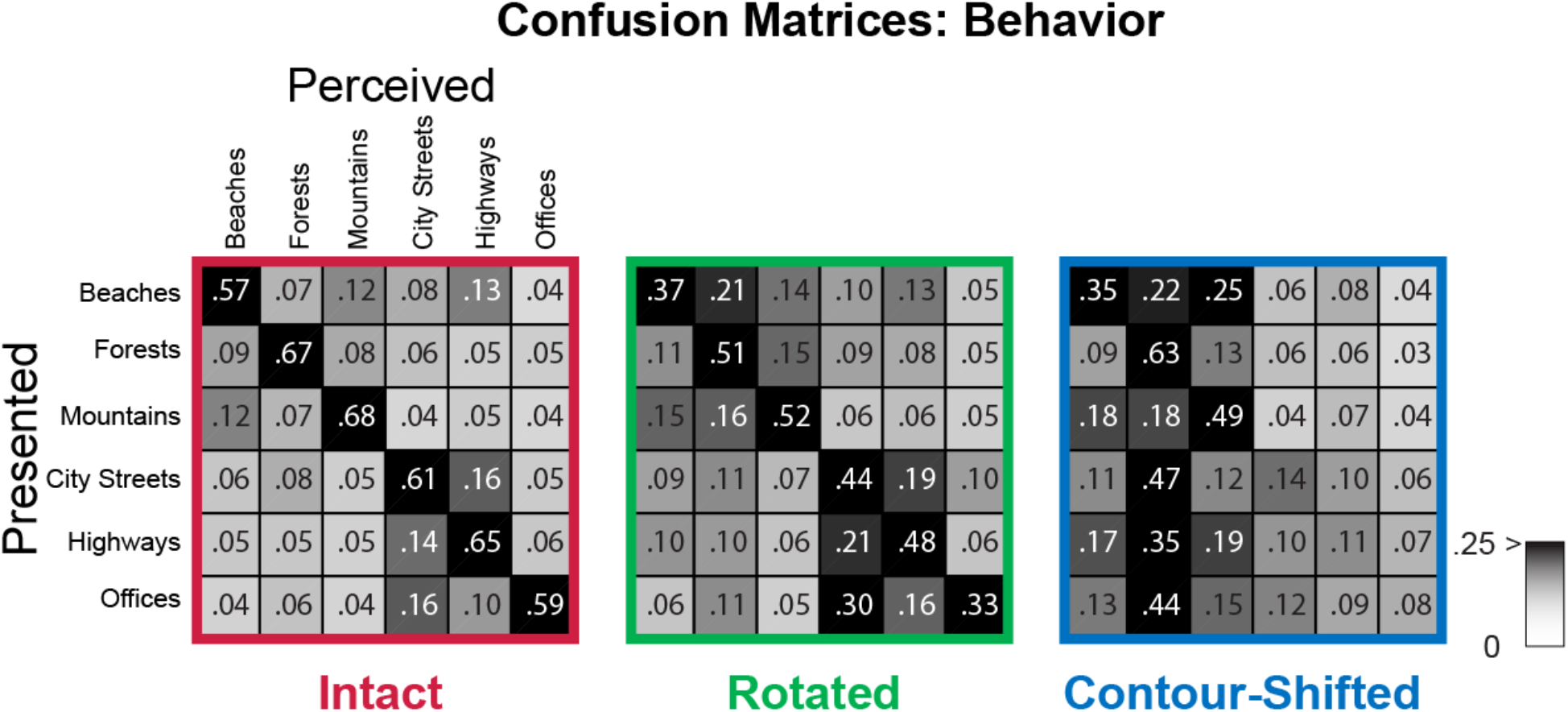
Group average confusion matrices of intact, rotated and contour-shifted line drawings obtained from a separate behavioral scene categorization experiment. The rows indicate true category labels presented to the participants, and columns indicate perceived category labels. In each entry, the probability of a perceived category given a presented category. The entries are also shaded according to the conditional probability: 0 as white, and 0.25 and higher as black.

**Figure S2.**
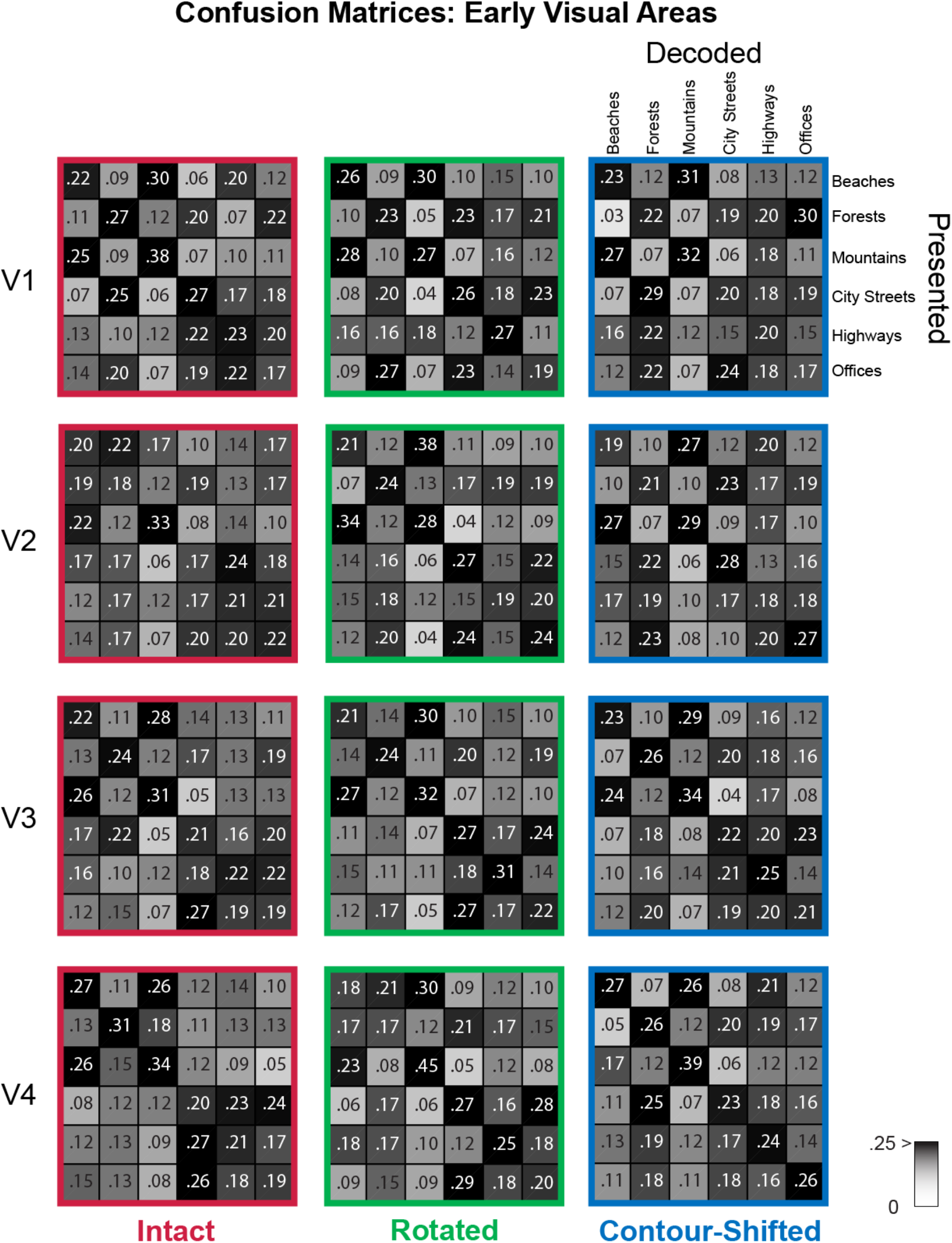
Group average confusion matrices of neural decoding from V1-4.

**Figure S3.**
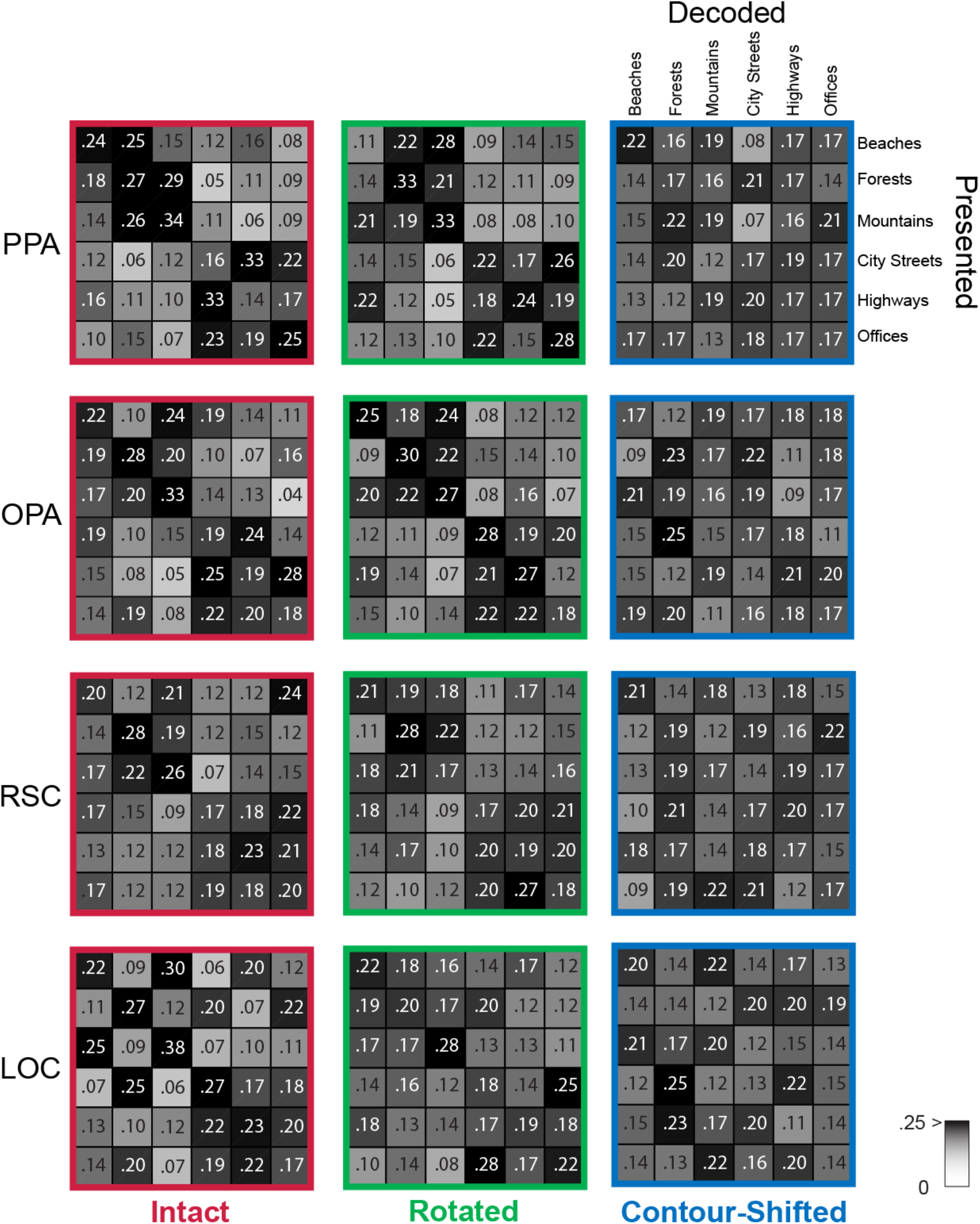
Group average confusion matrices of neural decoding from the PPA, OPA, RSC, and LOC.

### Un-thresholded error correlation and max property maps

**Figure S4.**
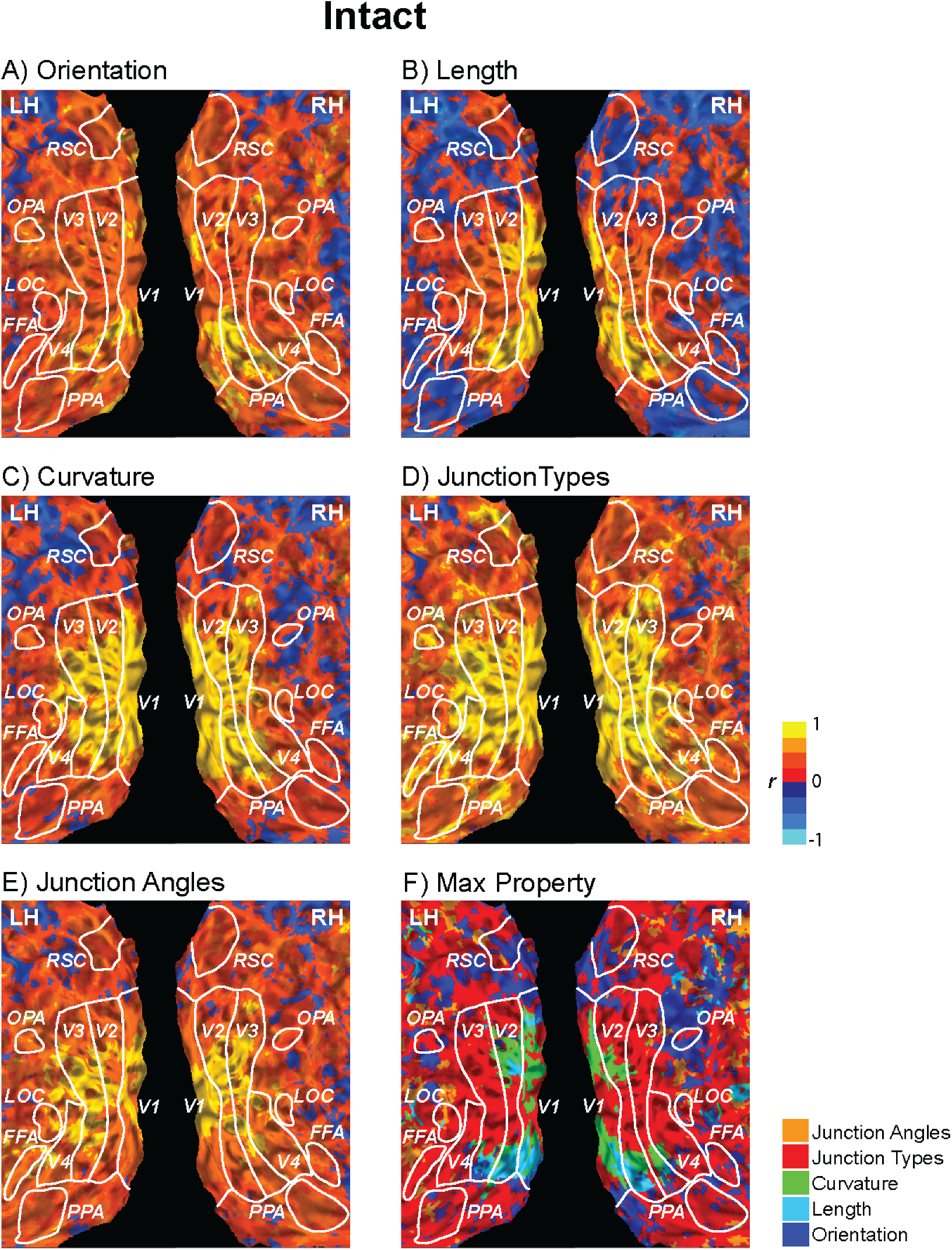
Un-thresholded error correlation maps between searchlight analysis of intact line drawings and computational scene categorization of the same intact line drawings. All searchlight locations were included regardless of their decoding accuracy. A-E) Searchlight locations are colored according to the strength of correlation between their neural decoding error patterns and computational error patterns (warm colors for positive and cold colors for negative correlation). F) Each searchlight location is colored according to the type of contour properties showing the maximum error correlation: orientation in dark blue, length in sky blue, curvature in green, junction types in red, and junction angles in orange.

**Figure S5.**
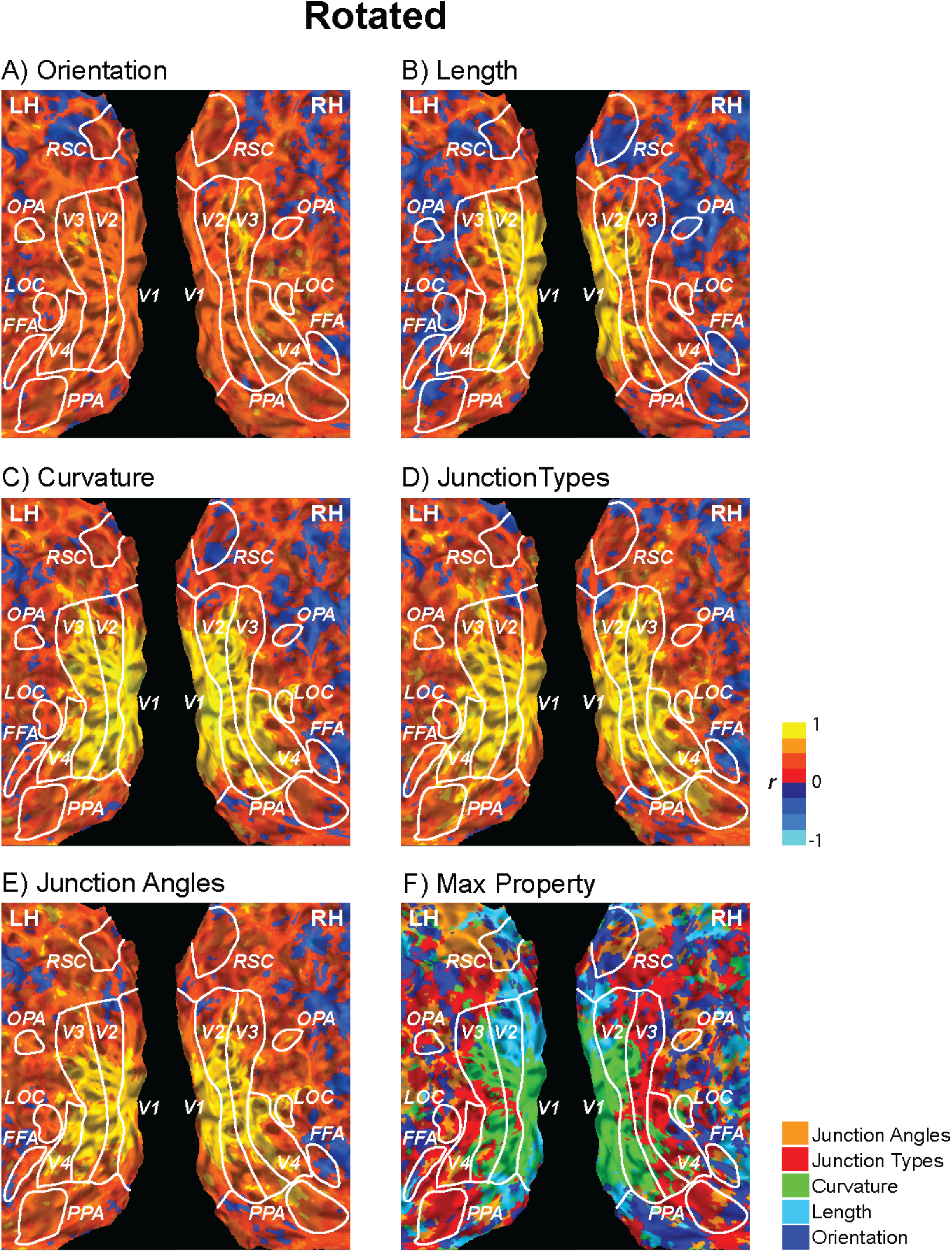
Un-thresholded error correlation and max property maps for rotated line drawings. Coloring follows the same conventions as Figure S4.

**Figure S6.**
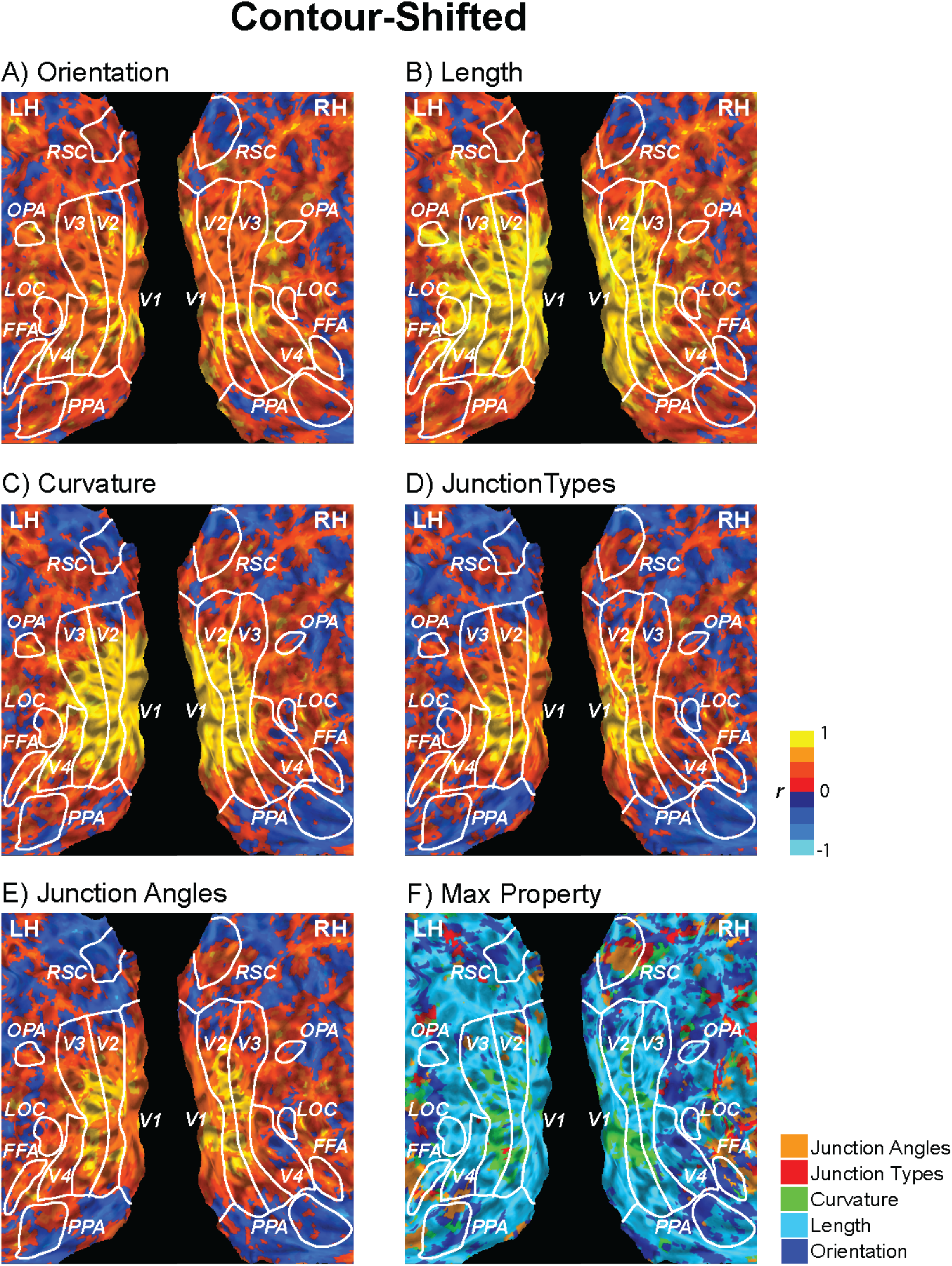
Un-thresholded error correlation and max property maps for contour-shifted line drawings. Coloring follows the same conventions as Figure S4.

### Error pattern correlation between brain and behavior: Searchlight analysis

We explored how disruption in contour orientation or junction properties affected the strength of error pattern correlation between neural decoding and behavior in a voxel-wise manner, using the same linear regression analysis as used for ROI-based analysis (see Methods in the main text). The linear mixed-effects modeling was performed with the fitlme function in MATLAB R2014b for faster computation speed compared to R. For each participant, the pattern of errors from the neural decoding analysis, i.e., the off-diagonal elements of the confusion matrix, was stored at each voxel location and registered to MNI space. Separately for each of the three line drawing types, neural decoding error patterns were Pearson-correlated to the group-averaged error patterns obtained from the separate behavioral experiment, resulting in 3-by-3 error correlation values for each voxel. Error correlations were transformed using Fisher’s z-transform. Using the same linear mixed-effects modeling shown in Figure 4C, we tested the extent to which the three idealized models can predict the three-by-three error correlation patterns between neural decoding and behavior. The coefficients were thresholded at *p* < .01 (two-tailed) with cluster correction of a minimum cluster size of 12 voxels. The three coefficient maps and their overlap are shown in Figure S7. Significant contributions from all three models (positive from same-type and intact-rotated, negative from intact-contour-shifted) are clearly discernible in both PPAs and the right RSC (red in Figure S7B). The overlap between the same-type and intact-rotated models is wide-spread throughout visual cortex (orange in Figure S7B). Significant contributions from all three models (positive from same-type and intact-rotated, negative from intact-contour-shifted) are clearly discernible in both PPAs and the right RSC (red in Figure S7B).

**Figure S7.**
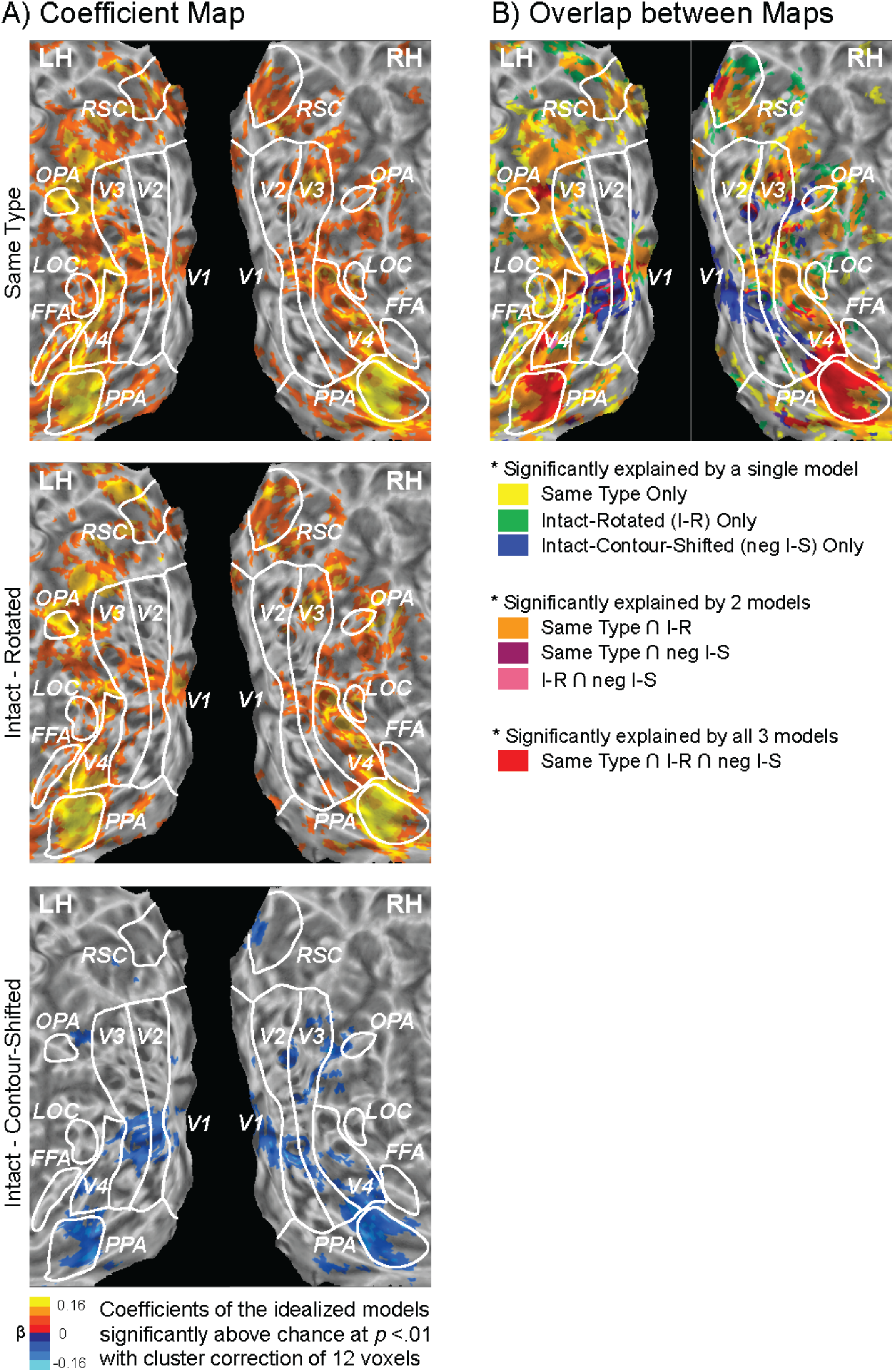
Searchlight analysis of patterns of error correlation between brain and behavior. At each searchlight locations the error correlation analysis were performed using the same three models used in the ROI analysis. A) The coefficient maps of the same type, intact-rotated, and intact-contour-shifted model. Each searchlight location is colored according to the coefficients, warm colors for positive and cold colors for negative. B) Each searchlight location is colored according to the set of models explaining patterns of error correlations at that location: only one model (yellow, green and blue), two of the models (orange, violet, and pink), or all three models (red). Consistent to the ROI results, the three idealized models significantly predicted

### Neural decoding using the equal number of participants across all ROIs

**Figure S8.**
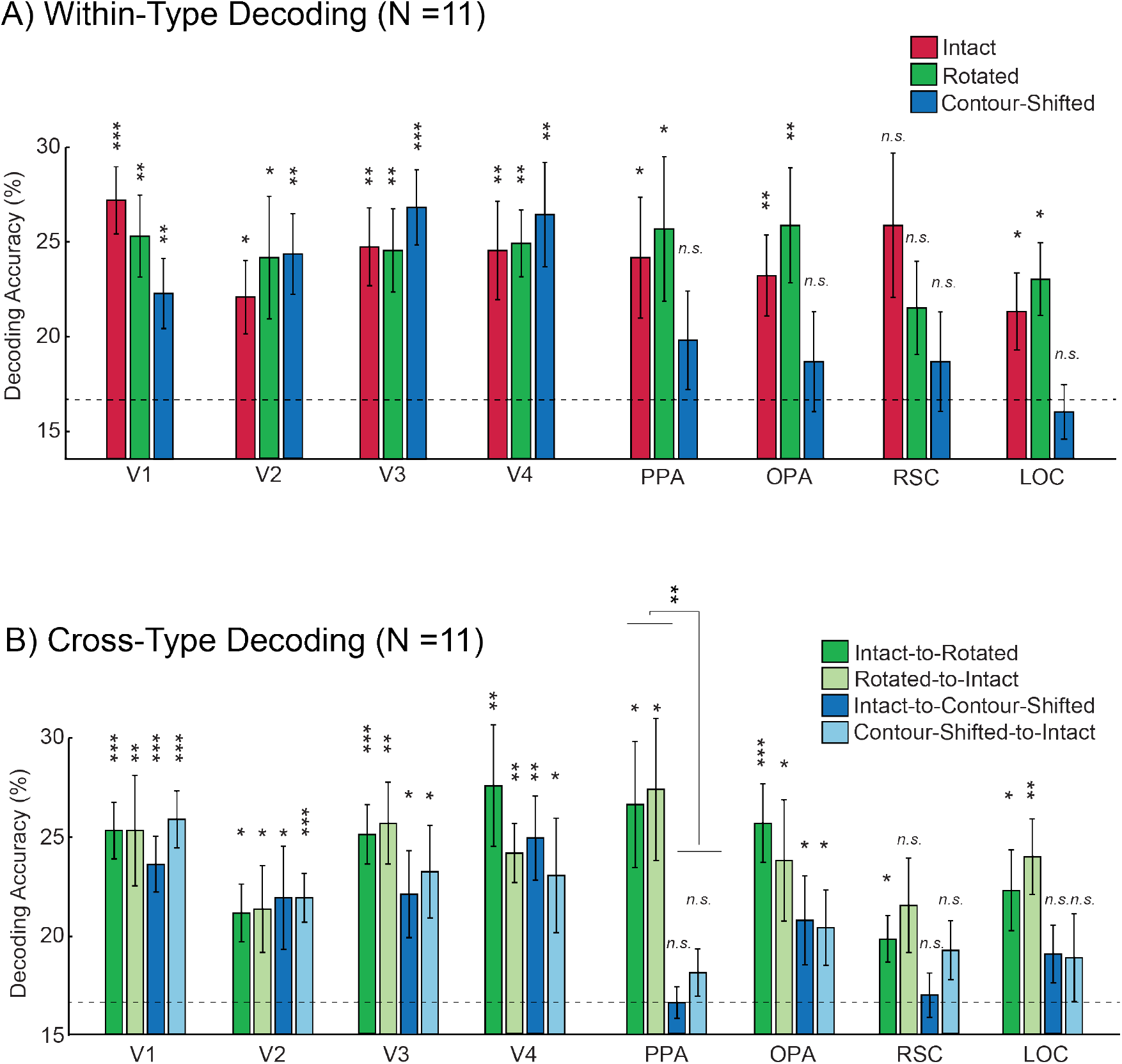
Neural decoding results from 11 participants in which we can delineate all nine ROIs (V1-4, PPA, OPA, RSC, LOC, and FFA). The patterns of results were identical to those from all 15 participants. A) Average accuracy rates of within-type category decoding from ROIs. C) Average accuracy rates of cross-type category decoding from the ROIs. The only significant difference in cross-type decoding accuracy between Intact-to-Rotated/Rotated-to-Intact and Intact-to-Contour-Shifted/Contour-Shifted-to-Intact was found in the PPA, *F*(1,10) = 12.154, *p* = .006, *η*^*2*^ = .549 as indicated above the bracket bridging the two bars. Error bars are standard errors of means. Dashed lines indicate chance performance (1/6). The significance of the one-sample t-test (one-tailed) was adjusted for multiple comparisons (FDR) and marked above each bar, \**q* < .05, \*\**q* < .01, \*\*\**q* < .001.

### Cross-type decoding between rotated and contour-shifted line drawings

**Figure S9.**
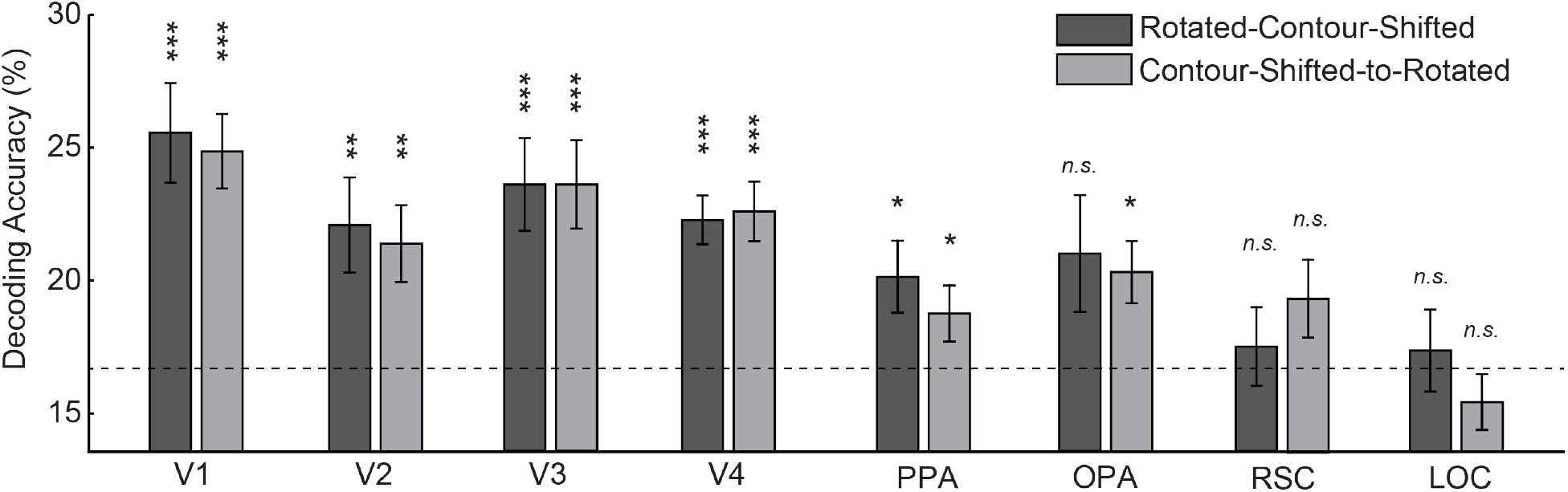
Average accuracy rates of cross-type category decoding between rotated and contour-shifted line drawings from the ROIs. Across the two types of line drawings, statistics of contour length and curvature are preserved, which is likely to underlie the robust cross-decoding accuracy in the early visual areas as well as in the OPA. By comparison, cross-decoding in the PPA is relatively weak. Error bars are standard errors of means. Dashed lines indicate chance performance (1/6). False discovery rate correction was employed to adjust significance for multiple comparisons. The significance of the one-sample t-test (one-tailed) is marked above each bar, \**q* < .05, \*\**q* < .01, \*\*\**q* < .001.

## Footnotes

1 2D-fast Fourier transform analysis of the triplets confirmed that our manipulation targeting the orientation statistics held in the Fourier space as well. Random contour-shifting had little impact on the Fourier amplitude spectrum, and the correlation in the Fourier amplitude spectrum were high between intact and rotated line drawings, Fisher’s z = 3.278 (*r* = .997). In contrast, the average correlation between the intact and rotated line drawings was relatively low, Fisher’s *z* = 1.475 (*r* = .901). The difference becomes even more pronounced when adjusting for the average correlation between images from different triplets; between intact and contour-shifted: *z*_*adj*_ = 1.073 (*r*_*adj*_= .791), between intact and rotated: *z*_*adj*_ = .030 (*r*_*adj*_ = .030).

2 Including these participants to the analysis did not change the pattern of results. However, as they acquire experience on the task, their relatively long presentation time reduced erroneous observations. Since our objective of having a brief presentation was to compare error patterns between neural decoding and behavior, we decided a priori not to include participants with SOAs < 100 ms to maximize the variance in error patterns.

